# Prediction of Cell States and Key Transcription Factors of the Human Cornea through Integrated Single-Cell Omics Analyses

**DOI:** 10.1101/2024.12.19.629354

**Authors:** Julian A. Arts, Sofia Fallo, Melanie S. Florencio, Jos G.A. Smits, Dulce Lima Cunha, Janou A.Y. Roubroeks, Mor Dickman, Vanessa L.S. LaPointe, Rosemary Yu, Huiqing Zhou

## Abstract

The cornea, a transparent tissue composed of multiple layers, allows light to enter the eye. Several single-cell RNA-seq analyses have been performed to explore the cell states and to understand the cellular composition of the human cornea. However, the inconsistences in cell state annotations between these studies complicate the application of these findings in corneal studies. To address this, we integrated single-cell RNA-seq data from four published studies and created a human corneal cell state meta-atlas. This meta-atlas was subsequently evaluated in two applications. First, we developed a machine learning pipeline cPredictor, using the human corneal cell state meta-atlas as input, to annotate corneal cell states. We demonstrated the accuracy of cPredictor and its ability to identify novel marker genes and rare cell states in the human cornea. Furthermore, cPredictor revealed the differences of the cell states between pluripotent stem cell-derived corneal organoids and the human cornea. Second, we integrated the single-cell RNA-seq based cell state meta-atlas with chromatin accessibility data, conducting motif-focused and gene regulatory network analyses. These approaches identified distinct transcription factors driving cell states of the human cornea. The novel marker genes and transcription factors were validated by immunohistochemistry. Overall, this study offers a reliable and accessible reference for profiling corneal cell states, which facilitates future research in cornea development, disease and regeneration.

**Significance statement:** This study creates a human corneal cell state meta-atlas that provides a common nomenclature of cells in the human cornea, through integrating multiple single-cell RNA-seq analyses. Using this meta-atlas, we developed a machine learning pipeline, cPredictor, to accurately annotate cell states in corneal studies using single-cell RNA-seq. Additionally, we identified distinct transcription factors driving cell states through integrating the atlas with chromatin accessibility data. This meta-atlas and the computational tool cPredictor enable future research in cornea development, disease, and regeneration.

## Introduction

The cornea, a multi-layered tissue, acts as a transparent protective shield located at the front of the eye. It allows light to enter the eye and functions as a focusing unit together with the lens. The tissue layers in the cornea have been extensively studied ^1,2^. The innermost layer of the cornea, the endothelium, is made of a thin layer of corneal endothelial cells. The endothelium maintains liquid homeostasis, important for maintaining correct corneal hydration^3^. The corneal stroma, composed mainly of keratocytes that produce extracellular matrix proteins, is anterior to the endothelium and plays a role in maintaining the transparency of the cornea as well as providing biomechanical strength for the cornea^4^. The corneal epithelium constitutes the outermost layer of the cornea. Composed of stratified epithelial cells linked together by desmosomes and tight-junctions, the corneal epithelium functions as a strong physical barrier^5^. Beyond the periphery of the cornea is the non-transparent conjunctiva. The conjunctiva has a role in providing ocular lubrication, as well as protecting the underlying tissues from insults, such as dust and microbes, entering deeper into the cornea^6^. Between the corneal epithelium and the non-transparent conjunctiva is the limbus. The limbus contains limbal stem cells that are capable of renewing the corneal epithelium and maintaining corneal regeneration^7–10^. The limbus also has a role as a physical barrier, preventing conjunctival cells and blood vessels from invading the transparent and avascular cornea^11^.

Various cell states in the different layers of the human cornea, including limbal, corneal epithelial, conjunctival, stromal, immune and vascular cells have recently been investigated by several studies using single-cell RNA-seq (scRNA-seq) analyses. Nevertheless, the number of defined cell states, their nomenclature and the corresponding marker genes do not agree among these studies, likely due to differences in data collection, processing and performed analyses^12–19^. The inconsistencies between these studies make it challenging to apply these findings in follow-up studies on corneal biology and disease. Therefore, integrating these studies to create a cell state meta-atlas that is comprehensive and can be used as a common reference^20^ is warranted. Additionally, most of these studies focus on identifying marker genes for cell states, whereas transcription factors (TFs) that play key roles in determining cell states were not centrally studied. Previous research on limbal stem cells reported a small number of TFs, such as PAX6, TP63, FOXC1, RUNX1, and SMAD3^21–25^. The key TFs in other corneal cell states remain less explored.

To reliably predict TFs controlling cell states, information on both gene expression (e.g., RNA-seq) and genomic regulatory elements that modulate gene expression needs to be incorporated. Regulatory elements are in accessible chromatin regions that can be detected by technologies such as ATAC-seq analysis^26^. TFs binding to the accessible regions can be predicted by motif analyses on the sequences in these genomic regions. RNA-seq and ATAC-seq data can further be integrated into gene regulatory networks (GRNs) that comprise TFs and their target genes (nodes), as well as the regulatory relationships between TFs and target genes (edges). By capturing the regulatory relationships, causality, and combinatorial interactions of TFs and their target genes, GRNs can enhance the ability to predict TFs that drive cell state differences. Thus, by combining gene expression and accessible chromatin regions detected at the single-cell level through scRNA-seq and scATAC-seq, and constructing single-cell GRNs, key TFs controlling specific cell states can be predicated in tissues containing heterogenous cell types such as the human cornea. Previously, we developed a computational pipeline, single-cell ANANSE (scANANSE)^27^, that combines scRNA-seq and scATAC-seq and generates pseudo bulk of these datasets to construct robust GRNs from these otherwise sparce data. scANANSE leverages TF expression, TF binding to anticipated target genes, and the expression of these target genes to build GRNs. scANANSE predicts the importance of a TF represented by an influence score through pairwise comparisons of GRNs of two cell states.

In this study, we integrated four publicly available scRNA-seq datasets of the human adult cornea to create a corneal cell state meta-atlas. This meta-atlas allowed us to annotate rare corneal cell states and identify novel marker genes, which were then experimentally validated. To facilitate its future application, we developed a support vector machine-based machine learning prediction pipeline, cPredictor, using this meta-atlas as the reference. As proof-of-principle, we showed that cPredictor could accurately annotate cell states in human corneal scRNA-seq studies on human adult corneas and on induced pluripotent stem cell (iPSC)-derived corneal organoids. Furthermore, integration of this scRNA-seq-based corneal cell state meta-atlas with the human cornea scATAC-seq data enabled us to construct GRNs using scANANSE and to identify key TFs driving various cell states in the human cornea.

## Results

### Data collection and integration of scRNA-seq of human corneal studies

To establish a cell state meta-atlas, we collected four scRNA-seq studies on adult human corneas from which data are currently publicly available^13–15,17^. Two of them were generated from complete corneas^13,14^. One study from Collin and colleagues^14^ used six donor corneas, one specifically for retrieving the limbal ring, one for central cornea and four complete corneas. Of note, in addition to scRNA-seq, this study also reported single-cell ATAC-seq (scATAC-seq) data that was applied later in this study for analyzing genomic regulatory regions. Another study, from Català and colleagues^13^, used eight donor corneas, and two of them specifically used the limbal ring. The 3^rd^ study, from Gautam and colleagues^15^, extracted cells from three whole eyes, of which the cornea data were included in this study. Additionally, we collected data from the study of Li and colleagues^17^ who isolated cells from the limbus of four donor corneas by removing the central cornea and superficial layers. In these four studies, different corneal sample disaggregation methods and data pipelines were used, which inevitably gave rise to different numbers and types of corneal cell states^28^.

To perform consistent analyses, we started with raw sequencing reads of all four studies. We performed data pre-processing, quality control and doublet removal with DoubletFinder^29^ on all datasets, and subsequently used scVI^30^ to integrate the data. scVI is currently considered one of the best integration tools for non-uniformly labeled datasets and is likely to maintain the biological relevance of cell states while minimizing batch effects^31^. Next, we applied unbiased Leiden clustering^32^ to group cells based on their gene expression profiles, identified highly variable genes of each group and annotated cell states with well-known markers among highly variable genes (Figure 1B). This clustering analysis resulted in a total of 21 distinct cell states, with a subdivision into two major branches: nine limbal/corneal epithelial-related clusters, and twelve non-epithelial diverse cell states including stromal cells and immune-related cells (Figure 1A). Notably, our data integration revealed a bias in cell retrieval across studies. For example, cells from cluster 3, limbal epithelial cells (LE), were mainly originated from the study of Li and colleagues^17^, clusters 8 and 9, central epithelial cells (CE), from Collin and colleagues^14^, and clusters 11 and 12, stromal keratocytes (SK) mainly from Catala and colleagues^13^ (Figure 1A).

**Figure 1:**
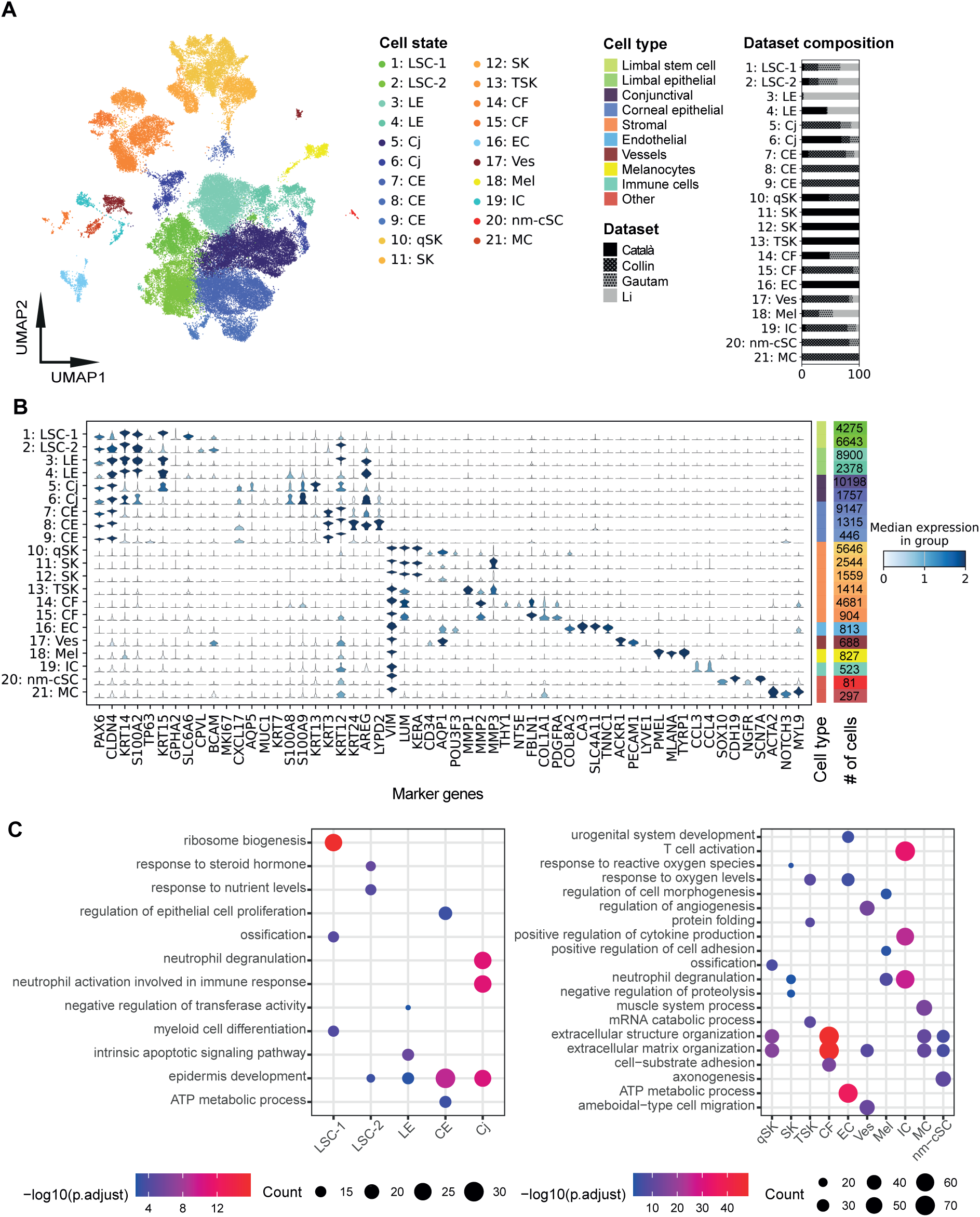
Integration, identification and characterization of cell states in the human cornea. **A)** UMAP of integrated and annotated cell states. Colors indicate overall cell type and sub colors define the respective cell states. The bar plots on the right depict dataset contributions for each cell state. **B)** Violin plot of literature annotated marker gene expression across integrated cell states. The height of the violin plot indicates the expression level of the markers and the color indicates the median score for all cells in that specific cell state. **C)** GO-term enrichment of selected cell states on all cluster markers identified with Seurat. Names of cell states (A): LSC-1 = Limbal stem cells 1; LSC-2 = Limbal stem cells 2; LE = Limbal epithelium; CE = Central epithelium; Cj = Conjunctiva; qSK =Quiescent stromal keratocytes; SK = Stromal keratocytes; TSK = Transitioning stromal keratocytes; CF = Corneal fibroblasts; EC = Endothelial cells; Ves = Vessels; Mel = Melanocytes; IC = Immune cells; nm-cSC = non-myelinating corneal Schwann cells; MC = Mural cells.

The branch of nine limbal/corneal epithelial-related clusters expressed *PAX6* (Figure 1B), a marker for limbal and corneal epithelial cells^33^ and *CLDN4*. This branch contained the largest number of cells, 45.059 out of 65.036 total. Although our nomenclature of cell states was apparently different from previous studies, our annotations of these cell states in the integrated meta-atlas were in general consistent with previous individual studies. For instance, corneal limbal cells from the study of Català and colleagues^13^ and limbal progenitor cells from Collin and colleagues^14^ were not annotated consistently in the two studies but clearly clustered together as a single limbal stem cell state in our cell state meta-atlas (Supplemental Figure 1 and 2).

The branch represented by twelve non-epithelial cell states expressed *VIM* and consisted of non-epithelial cells, such as stromal cells, fibroblasts and immune cells. The immune related cluster was consistent with immune cells from previous studies^14,15^. However, cells in stroma were annotated differently in previous studies. Specifically, cells classified as fibroblasts in the study by Gautam and colleagues^15^ and stromal keratocytes in the study of Català and colleagues^13^ grouped together, and they were indistinguishable as corneal fibroblasts (CF) in our meta-atlas.

Likewise, corneal stromal cells from Collin and colleagues^14^ and stromal keratocytes from Català and colleagues^13^ clustered together and were annotated as quiescent stromal keratocytes (qSK). Notably, we identified a previously unidentified cell state showing distinct markers for non-myelinating corneal Schwann Cells (nm-cSC). They contained a small number of cells (<5%), previously identified as fibroblasts, melanocytes and corneal endothelial cells from individual studies (Supplemental Figure 1 and 2). These cells were not identified as nm-cSCs, likely due to their low number in individual studies, highlighting the power of integrating the studies.

### Distinct and shared marker genes in the cell state meta-atlas of the human cornea

In the limbal/corneal epithelial branch, clusters 1-4 were identified as limbal cell states based on the expression of stem cell and limbal marker genes. Cluster 1 and cluster 2 both expressed limbal markers *CXCL14*^34^ and *KRT14*^35^ and the stem cell marker *TP63*, and were therefore annotated as limbal stem cells (LSC) (Figure 1B). PROGENy pathway analysis^36^ of highly variable genes (HVGs) revealed WNT pathway enrichment in both clusters (Supplemental Figure 3). Among these two clusters, cluster 1 (LSC-1) highly expressed *KRT15*^37^ and *GPHA2*. *GPHA2* has been associated with early/quiescent LSC state^38,39^. In cluster 1, we also identified a novel marker gene *SLC6A6*. Cluster 2 (LSC-2) expressed *CPVL* that was reported as a limbal stem cell niche marker^13^. We confirmed marker gene expression at the protein level by immunochemical staining. SLC6A6 and CPVL were both expressed in the limbal region and showed small overlap with p63 (Figure 2A and 2B). LSC-2 also expressed differentiation-related genes like *KRT3* at a low level, suggesting that LSC-2 represents an early stage of differentiated limbal stem cells^40^. Clusters 3 and 4 both expressed the limbal marker *KRT14*, sharing GO-term^41^ enrichment for epidermal development (Figure 1C) and the EGFR pathway (Supplemental Figure 3), but had low *TP63* and *KRT15* expression. They were annotated as limbal epithelium (LE). These two clusters differed slightly in the marker gene expression, *CXCL14* and *GJB6* higher in cluster 3, and *S100A2* and *AREG* higher in cluster 4.

**Figure 2:**
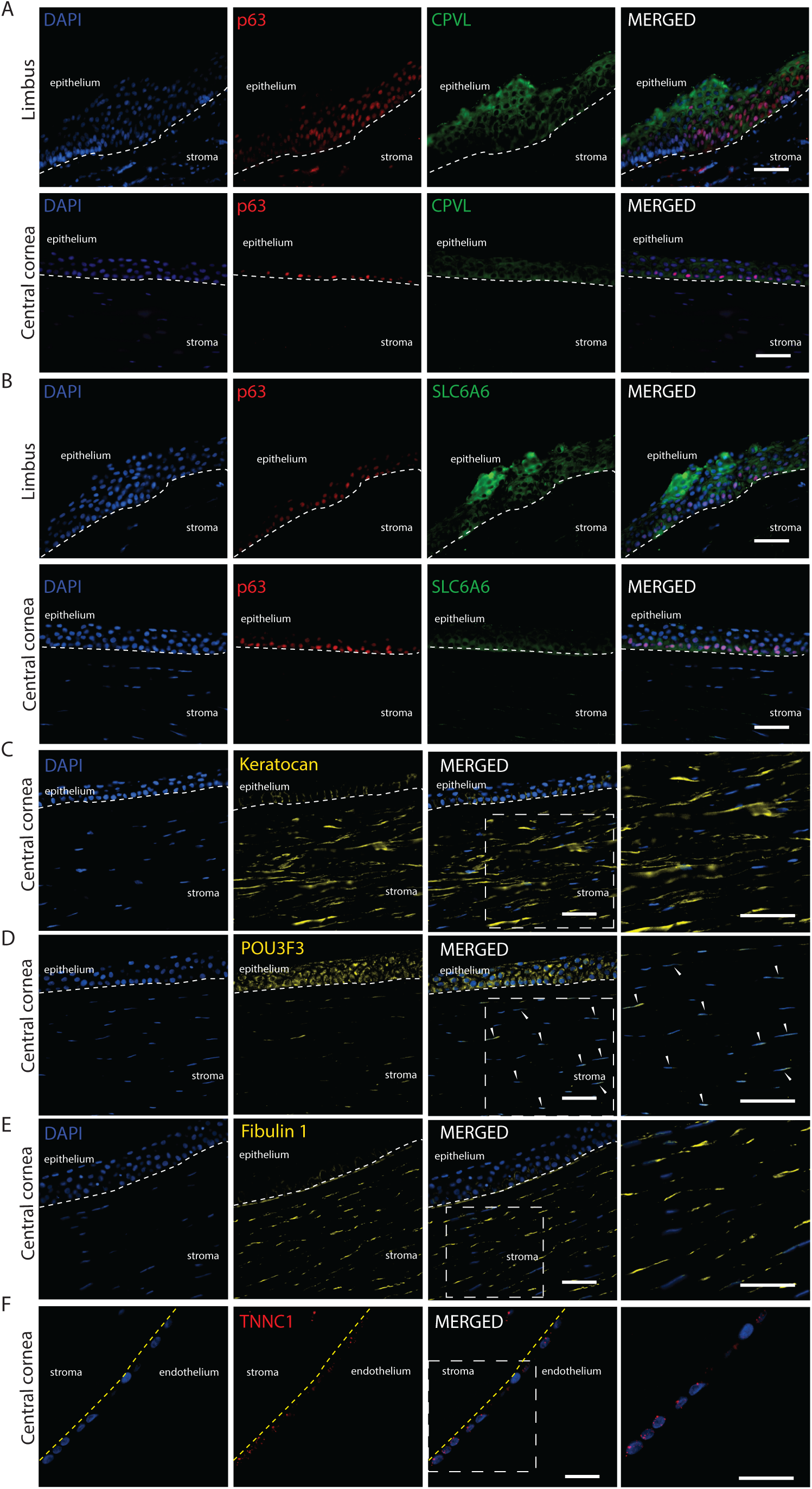
Immunofluorescence of novel markers in the human cornea. **A-B)** Co-staining of CPVL and SLC6A6 (green) with p63 (ΔNp63) (red) in limbus and central cornea. **C-E)** Staining of corneal stroma marker Keratocan (C), POU3F3 protein (D) and Fibulin-1 (E) (yellow). **F)** Staining of TNNC1 (red) in corneal endothelium. Arrowheads depict cells in the stroma with nuclear expression of POU3F3. Dotted lines indicate borders between stroma and epithelium (in white) and between stroma and endothelium (in yellow). Cell nuclei were stained with DAPI (blue). Scale bar represents 50 μm.

Other clusters 5–9 in the limbal/corneal epithelial branch all expressed the epithelial mucosal barrier marker *CXCL17*^42^(Figure 1B) and HVGs were strongly enriched for the epidermis development (GO) (Figure 1C) and for the p53 pathway (Supplemental Figure 3). Clusters 5 and 6 were annotated as conjunctiva (Cj) cells, marked by expression of *AQP5*^43^, *MUC1*^44,45^, *KRT7*^46^, *KRT13*^47^ and *S100A9*^48^. Pathway analysis of these conjunctiva clusters showed their strong link to immune responses, such as neutrophil degranulation, neutrophil activation, and the TNF-α pathway^49,50^ (Supplemental Figure 3). Clusters 7-9 were annotated as central epithelium (CE) cells, as they all expressed *KRT3* and *KRT12*^40,51^. Clusters 7 and 8 had higher *KRT24*^52^ and *LYPD2*^14^ expression, typically for central epithelial cells, whereas cells from cluster 9 had highest *KRT3*, *KRT12* and *PAX6* expression, suggesting the most differentiated corneal epithelial cells in this cluster.

In the non-epithelial branch, clusters 10–15 were identified as stromal cells due to their expression of the corneal stroma marker *LUM*^53^ (Figure 1B). The clusters separated into multiple distinct cell states and were annotated as stromal keratocyte and fibroblast clusters. Among these clusters, cluster 10 was identified as quiescent stromal keratocytes (qSK) due to its high expression of *LUM* and *KERA*^53^ as well as quiescent stromal keratocyte markers *CD34*^54^ and *AQP1*^55^. Notably, cells from this cluster expressed high levels of the transcription factor *POU3F3*, a novel marker gene for qSK. By co-staining stromal cells with the keratocyte marker Keratocan (encoded by *KERA*) (Figure 2C), we confirmed that POU domain, class 3 (POU3F3 protein, encoded by *POU3F3)* expression was localized in the nuclei of a small subset of stromal cells (Figure 2D), consistent with its mRNA-level expression detected by scRNA-seq in qSK. In contrast to the nuclear expression of *POU3F3* at both mRNA and protein levels in stromal cells, POU3F3 protein was also detected in the cytoplasm of central corneal basal cells where no *POU3F3* mRNA was detected. Clusters 11 and 12 were annotated as stromal keratocytes (SK) as they displayed lower *CD34* expression but high levels of the stromal keratocyte marker *KERA*^56^.

Cluster 13 cells expressed *MMP1* and *MMP3*, known for their involvement in stromal keratocyte extracellular matrix remodeling and mechanical stress responses^18,57,58^, which are critical processes in stromal keratocytes that transition into a repair-like phenotype^59^. We therefore annotated this cluster as transitional stromal keratocytes (TSK). Clusters 14 and 15 were defined as corneal fibroblasts (CF) because cells in these clusters exhibited high expression of general fibroblast markers *FBLN1*^60^, *COL1A1*, and *COL5A1*^61^ (Figure 1B). We confirmed the expression of the fibroblast marker Fibulin 1 (encoded by *FBLN1*) in a subset of stromal cells, likely to be fibroblasts (Figure 2E). GO and pathway analyses showed that qSK and fibroblast clusters (10, 14, 15) were enriched in genes involved in extracellular matrix and structure organization (Figure 1C), and in Androgen, Estrogen, TNF-α and PI3K signaling pathways (Supplemental Figure 3), known to be involved in stroma cell function^62,63^.

Clusters 16–21 within the non-epithelial branch displayed non-stromal identities and had a relatively small number of cells. Cluster 16 was identified as endothelial cells (EC) due to its unique expression of *CA3*, *COL8A2* and *SLC4A11*^64–66^ (Figure 1B). Interestingly, cells in this endothelial cluster expressed high levels of *TNNC1*, which was validated by protein staining of Troponin C1 (Figure 2F). Although the function of Troponin C1 in corneal endothelial cells is not yet clear, *TNNC1* may be a novel marker for this cell state. Furthermore, cluster 16 exhibited enrichment in GO terms related to ATP metabolism, oxygen level regulation (Figure 1C) and the hypoxia pathway (Supplemental Figure 3), consistent with the known function of endothelial cells in fluid pumping that requires energy^67,68^. Cluster 17 was annotated as vessels (Ves) since most cells uniquely expressed the blood vessel marker *ACKR1* (Figure 1B). Additionally, this cluster expressed the lymph vessel marker *LYVE1*^69,70^ and displayed enrichment in the vascular endothelial growth factor (VEGF) pathway (Supplemental Figure 3). Cluster 18 was annotated as melanocytes (Mel) due to its high expression of the melanocyte markers *PMEL*^71^, *MLANA*^72^ and *TYRP1*^73^ (Figure 1B). Cluster 19 was identified as immune cells (IC) because its cells specifically expressed *CCL3 and CCL4*, markers for T-cells and macrophages^74^ (Figure 1B) and demonstrated significant enrichment in functions such as T-cell activation, positive regulation of cytokine production (Figure 1C), and the NF-κB pathway (Supplemental Figure 3). The smallest cluster 20 was annotated as non-myelinating corneal Schwann cells (nm-cSC) as its cells exclusively expressed Schwann cell markers such as *SOX10*^75^*, CDH19*, *NGFR*^76^, and the non-myelinating corneal Schwann cell marker *SCN7A*^77^ (Figure 1B). Lastly, cluster 21, a cluster previously defined as Fibroblast Corneal Endothelial Cells by Collin and colleagues^14^ (Supplemental Figure 1 and 2), was annotated as mural cells (MC) since cells of this cluster expressed unique mural cell markers *ACTA2*, *NOTCH3* and *MYL9*^61,78^ (Figure 1B).

In summary, we created a corneal cell state meta-atlas that contains more comprehensive annotations of corneal cell states and their associated marker genes, through integrating multiple scRNA-seq analyses.

### Developing a machine learning-based prediction tool for human corneal cell states using the meta-atlas as input

To facilitate corneal cell state analysis in future scRNA-seq studies, we constructed cPredictor, a machine learning pipeline that leveraged cell state annotations of our meta-atlas to train a support vector machine (SVM) model. An SVM was selected since this model works well for scRNA-seq dataset annotations^79^ and it enables straightforward model explainability. To do this, we first performed four rounds of recursive feature elimination (RFE) using SHapley Additive exPlanations (SHAP^80^), reducing the feature space to 1243 genes (consisting of 1047 HVGs and 196 non-HVGs; Supplemental Figure 4A) that retained known marker genes of corneal cell states, such as *PAX6, KERA, FLBN1, SCL4A11, SCN7A* and *NOTCH3* (Figure 1B). We then conducted hyperparameter tuning on the regularization parameter, class weights and number of iterations (Material and Methods) and performed a 5-fold cross validation with the 1243 selected genes. This resulted in our final model hyperparameters: 0.01 for the regularization parameter, balanced class weights and 1000 maximum number of iterations, with a model performance of a weighted F1-score of 95.75% across all classes (Figure 3B). Model calibration was performed to ensure that the model-predicted cell state certainty scores closely resembled the observed probabilities during model training. In addition to cell state prediction, cPredictor also outputs common machine-learning scores and calibration plots, indicating both the performance of the trained model and calibration for each class. Moreover, cPredictor generates pretrained models from which the top n genes driving the cell state predictions (top explainable genes) can be investigated by Explainable AI methods, such as SHAP.

**Figure 3:**
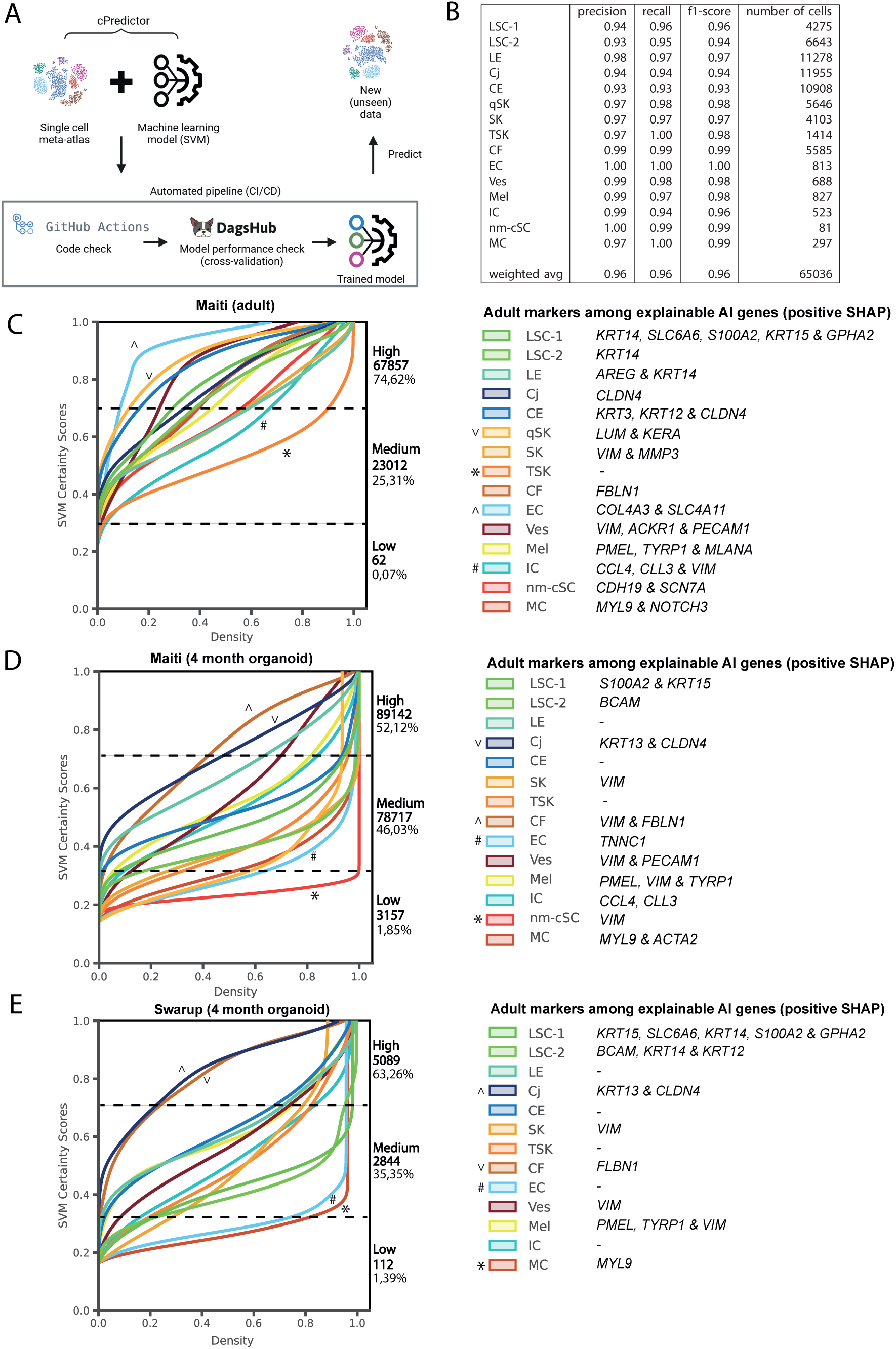
A machine learning-based prediction pipeline, cPredictor, for human corneal cell states in scRNA-seq datasets. **A)** Schematic overview of cPredictor’s machine learning pipeline. cPredictor applies the human corneal meta-atlas as reference data to support vector machines (SVM), which enables annotation of adult corneal cell states in external datasets. **B)** Sklearn’s^82^ classification report on the performance of a five-fold cross-validation using the cornea cell state meta-atlas. **C-E)** Prediction certainty plots of corneal cell states on adult corneal cells (C), 4-month-old organoids from Maiti^19^ (D), and 4-month-old organoids from Swarup^81^ (E). Left, the x-axis shows the cumulative kernel densities, and the y-axis depicts the model confidence (SVM certainty score). The numbers and percentages of cells corresponding to low (< 0.3), medium (>0.3 and <0.7) and high (> 0.7) certainty scores in each dataset are depicted next to the plots. ^/V indicate cell states most similar and */# indicate cell states least similar to corneal cell states from the meta-atlas. Right, corneal markers among the top ten explainable AI genes driving model’s decisions for each of the predicted corneal cell states determined by their positive SHAP values are shown.

To test the performance of cPredictor, we applied it to predict the corneal cell states on one dataset from human adult cornea^19^ that was not used during model training and cross-validation. This resulted in highly confident cell state annotations (certainty scores >0.7 on a scale of 0-1) for most cells (∼75%; Figure 3C). The remaining cells (∼25%) had medium certainty scores (>0.3 and <0.7), and only 62 cells (<0.1%) showed low certainty scores (<0.3). Compared to that original study^19^ in which twelve clusters were identified, cPredictor identified all fifteen clusters with high or medium certainty scores. Among these clusters, endothelial cells (EC) and quiescent stromal keratocytes (qSK) had highest certainty scores (>0.7), immune cells (IC) and transitioning stromal keratocytes (TSK) had lowest scores (<0.3) compared to all other cell states. Different limbal and corneal epithelial cell states showed high and medium certainty scores, accompanied with known marker genes (Figure 1B) among the top ten explainable genes (Supplemental Figure 5A). These included *KRT14*, *SLC6A6* and *S100A2, KRT15* and *GPHA2* for LSC-1, *KRT14* for LSC-2, *AREG* and *KRT14* for LE, *KRT3*, *KRT12* and *CLDN4* for CE, and CLDN4 for Cj. Cells from non-epithelial cell states cells also showed high and medium certainty scores with known markers among the top explainable genes, such as *LUM* and *KERA* for qSK, *VIM* and *MMP3* for SK, *FBLN1* for CF, *COL4A3* and *SLC4A11* for EC, *VIM*, *ACKR1* and *PECAM1* for Ves, *PMEL*, *TYRP1* and *MLANA* for Mel, *CCL3*, *CCL4 and VIM* for IC, *MYL9 and NOTCH3* for MC and *CDH19* and *SCN7A* for nm-cSC. It is worth noting that, in this dataset, cPredictor also annotated 196 cells as MC (0,2%) and 101 cells (0,1%) as nm-cSC that were not identified in the original study, likely due to their small number.

To summarize, we demonstrated that cPredictor was able to predict cell states including the rare types on a scRNA-seq dataset of the human cornea.

### Assessing cell states in induced pluripotent stem cell derived corneal organoids using cPredictor

Corneal organoids derived from induced pluripotent stem cells (iPSC) represent good models to study corneal biology and pathogenesis. However, how cell states in iPSC-derived corneal organoids resemble those in human cornea is still an open question. To address this, we collected two studies where scRNA-seq was performed on iPSC-derived corneal organoids; one from Maiti and colleagues^19^ using 4-month-old organoids, and the other from Swarup and colleagues^81^ containing time series data from 1-month-old organoids up until 4-month-old organoids.

First, we applied cPredictor to the two scRNA-seq datasets of 4-month-old organoids^19,81^ to determine cell states, as the 4-month-old organoids may be most similar to the corneal cell state meta-atlas that was created using data from human adult corneas. In the study of Maiti and colleagues^19^, cPredictor annotated ∼52% of the cells with high certainty scores (>0.7) and ∼46% with medium certainty scores (>0.3 and <0.7). There was a small number of cells annotated with low certainty scores (∼2%; Figure 3D). Limbal/corneal-epithelial cell states mainly showed medium certainty scores. Compared to cell states in the meta-atlas (Figure 3C), there was a reduction in the number of markers among the top explainable genes (Figure 3D and Supplemental Figure 5B). These included *SLC6A6* and *KRT15 for LSC-1*, *BCAM* for LSC-2 and no well-known markers for LE and CE. Cj showed highest certainty scores (>0.7) among epithelial cell states, together with *KRT13* and *CLDN4* among the top explainable genes. Most non-epithelial cells showed medium certainty scores, with a reduced number of markers among the top explainable genes (Figure 3D and Supplemental Figure 5B). This included *VIM* for the majority of cell states, *TNNC1* specifically for endothelial cells (EC), *PMEL* and *TYRP1* for Melanocytes (Mel), *CCL3* and *CLL4* for immune cells (IC), *MYL9* and *ACTA2* for mural cells (MC). Among them, CF showed high certainty scores (>0.7) together with *VIM* and *FBLN1* among the top explainable genes. Moreover, nm-sSC showed low certainty scores (<0.3) without *CDH19* and *SCN7A* among the top explainable genes, and no cells were predicted as qSKs.

In the other study where 4-month organoids were also generated^81^, cPredictor showed similar annotation of cell states, ∼63% showed high, ∼35% showed medium and ∼1% showed low certainty scores (Figure 3E). Similar to the data from Maiti and colleagues^19^, Cj and CF cells showed highest certainty scores (>0.7), contributing to the majority of cells having highest certainty scores, and none of the cells were predicted as qSK. LSC-1 and LSC-2 also showed similar certainty scores, as compared to those from Maiti and colleagues, but with additional markers in the top explainable genes, *KRT14* and *GPHA2* for LSC-1 and *KRT12* and *KRT14* for LSC-2. Other differences included mural cells (MC), which showed very low certainty scores (<0.3), with only *MYL9* among the top explainable genes, and no nm-cSC cells were predicted. As expected, cell states from earlier timepoints of organoids (month 1-3) showed even less cells having high certainty scores (>0.7): 29% at 1 month, 36% at 2 months, and 37% at 3 months (Supplemental Figure 4B-D), with a limited number of well-known markers among the top explainable genes (Supplemental Figure 5D-F).

Taken together, our cPredictor results showed a large difference between cell states of iPSC-derived corneal organoids and those of the human adult corneas, suggesting more immature cell states in corneal organoids.

### Integrating human cornea scATAC-seq data with the corneal cell state meta-atlas

Having obtained comprehensive cell states in the human cornea, we sought to identify key regulators that govern cell state determination as the second application of the human corneal cell state meta-atlas. For this, we used scATAC-seq data of the human cornea^14^. To combine this dataset with the scRNA-seq based corneal cell state meta-atlas, we employed the Seurat label transfer method to label the cell state of each single cell in the scATAC-seq data. This method matches DNA accessibility signals at genomic regions near expressed genes detected in scRNA-seq, and predicts and labels scATAC-seq cells according to the cell states in scRNA-seq. In addition, the confidence of the prediction is represented by a model prediction score. Using this method, we successfully identified all cell states that were defined in the meta-atlas in the scATAC-seq, except MC (Figure 4A). Since scATAC-seq data was known to be sparce, we selected cells with a model prediction score of 0.4 or higher (range 0-1) and clusters consisting of at least 100 cells as reliable data for downstream analyses. This gave rise to cell clusters LSC-1, LSC-2, LE, CE, Cj, qSK and CF (Figure 4B). Among limbal/corneal epithelial cell states, LSC-1 appeared to be clearly distinct from others, whereas other limbal/corneal epithelial cells (LSC-2 and LE, CE, Cj) clustered together, based on scATAC-seq. Predicted Cj cells had the highest contribution among cell states in scATAC-seq. qSK and CF were distinct from the limbal/corneal epithelial cells and separated into small clusters (Figure 4B).

**Figure 4:**
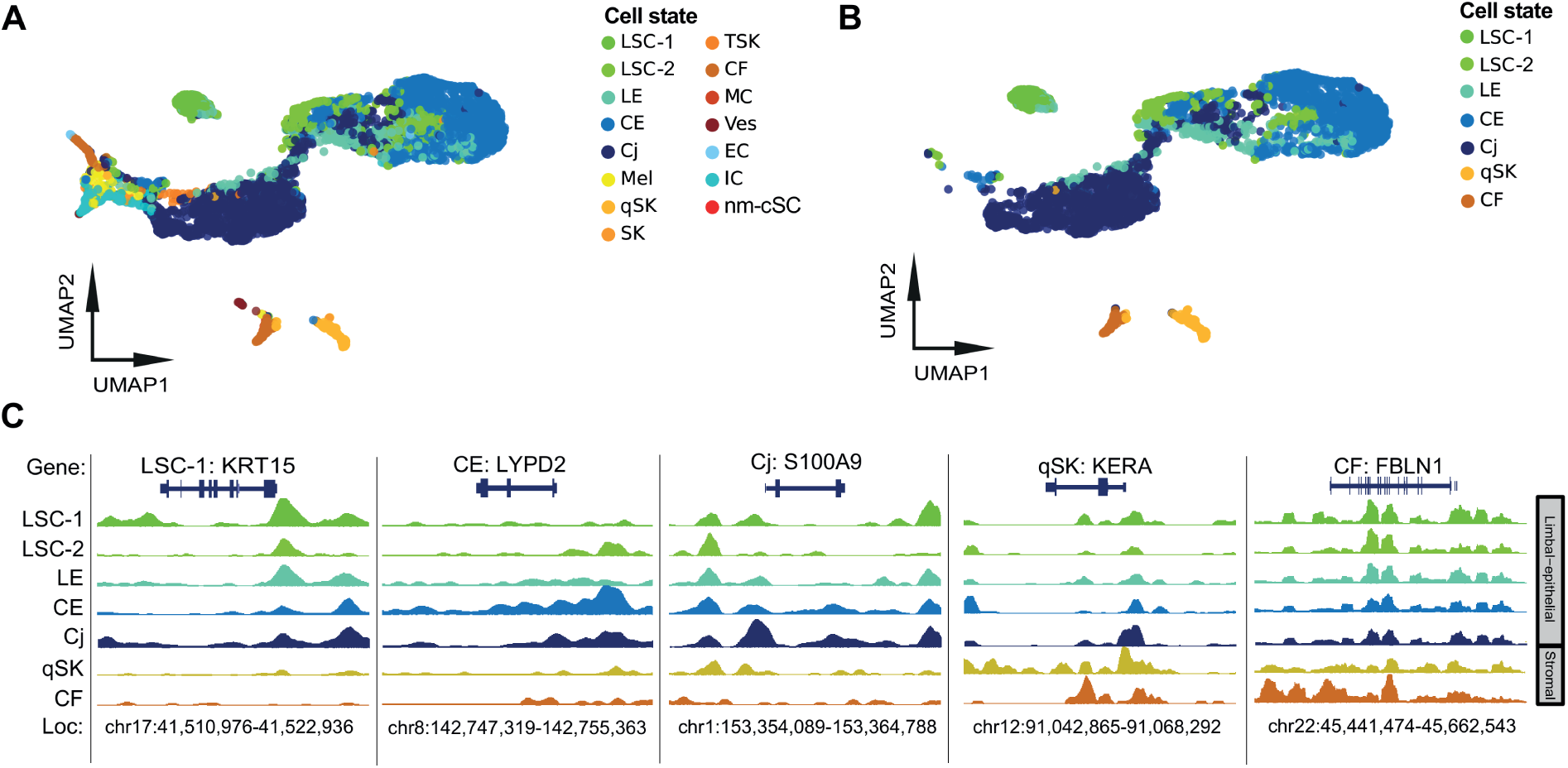
Characterization of scATAC-seq signals across cell states in the human cornea. **A)** UMAP of scATAC-seq imputed from scRNA-seq using Seurat label transfer. **B)** UMAP of sub selected cell states with at least 100 cells and a minimal prediction score of 0.4. **C)** Genome browser tracks showing open chromatin profiles around cell state specific marker genes. Names of cell states: LSC-1 = Limbal stem cells 1; LSC-2 = Limbal stem cells 2; LE = Limbal epithelium; CE = Central epithelium; Cj = Conjunctiva; Mel = Melanocytes EC = Endothelial cells; IC = Immune cells; Ves = Vessels; qSK = Quiescent stromal keratocytes; SK = Stromal keratocytes; TSK = Transitioning stromal keratocytes; CF = Corneal fibroblasts; nm-cSC = non-myelinating corneal Schwann cells

To confirm the predicted cell states in scATAC-seq through label transfer, we examined scATAC-seq signals near a select subset of marker genes from scRNA-seq (Figure 1B). Among limbal/corneal epithelial cell states, the scATAC-seq signal of the marker gene *KRT15* was highest in LSC-1, consistent with its expression in scRNA-seq (Figure 1B), indicating appropriate label transfer prediction for this cell state (Figure 4C). *LYPD2*, an identified marker gene for CE displayed a high scATAC-seq signal in CE (Figure 4C). As expected, Cj cells displayed the highest accessibility for *S100A9*, a conjunctival marker gene (Figure 4C). In the stromal cell states, the highest accessibility of the marker gene *KERA* was observed in qSK, as compared to other cell states (Figure 4C). Similarly, the fibroblast marker *FBLN1* exhibited high accessibility in CF (Figure 4C). These results indicated that the label transfer method gave reasonably accurate prediction of cell states using scATAC-seq data, but also showed less distinct separation of cell states, as compared to using scRNA-seq.

### Prediction of key transcription factor using motif analysis on accessible chromatin regions

Next, to predict binding of key transcription factors (TF) driving cell states, we generated pseudo bulk for each cell state by merging cells from cell states, and subsequently performed TF motif enrichment analysis on accessible chromatin regions detected by scATAC-seq. We then compared the TF motif enrichment scores with pseudo bulk gene expression of linked TFs through correlation analysis. This identified expressed TFs binding to accessible chromatin regions and potentially highly important in driving gene expression in specific cell states.

We identified two distinct groups of TFs associated with either the limbal/corneal epithelial or the non-epithelial branch. The group associated with the limbal/corneal epithelial cell states included many known TFs in corneal limbal stem cells or epithelial cells such as TP63, FOSL2, PAX6, GRHL1, OTX1, SMAD3 and RXRA (Figure 5A). Among these TFs, TP63 showed most specific high scATAC-seq signals at the TP63 motif in all three limbal cell states, LSC-1, LSC-2 and LE (Figure 5A and 5B), consistent with highest expression in LSC-1. SMAD3 displayed a high TF binding enrichment score only in LSC-2, despite its broader expression (Figure 5A). FOSL2, PAX6 and GRHL1 showed high TF binding motif enrichment scores in several limbal and epithelial cell states (Figure 5B), consistent with their relatively high gene expression in these cell states (Figure 5A and 5B). OTX1 had high TF binding motif enrichment scores in both LSC-1 and Cj, in line with its gene expression in these cell states. RXRA that was highly expressed in CE showed high TF binding enrichment score unique to CE (Figure 5A). Interestingly, ZEB1 exhibited an inverse relationship between TF binding motif enrichment and gene expression, suggesting a repressor role of this TF in limbal/corneal epithelial cells.

**Figure 5:**
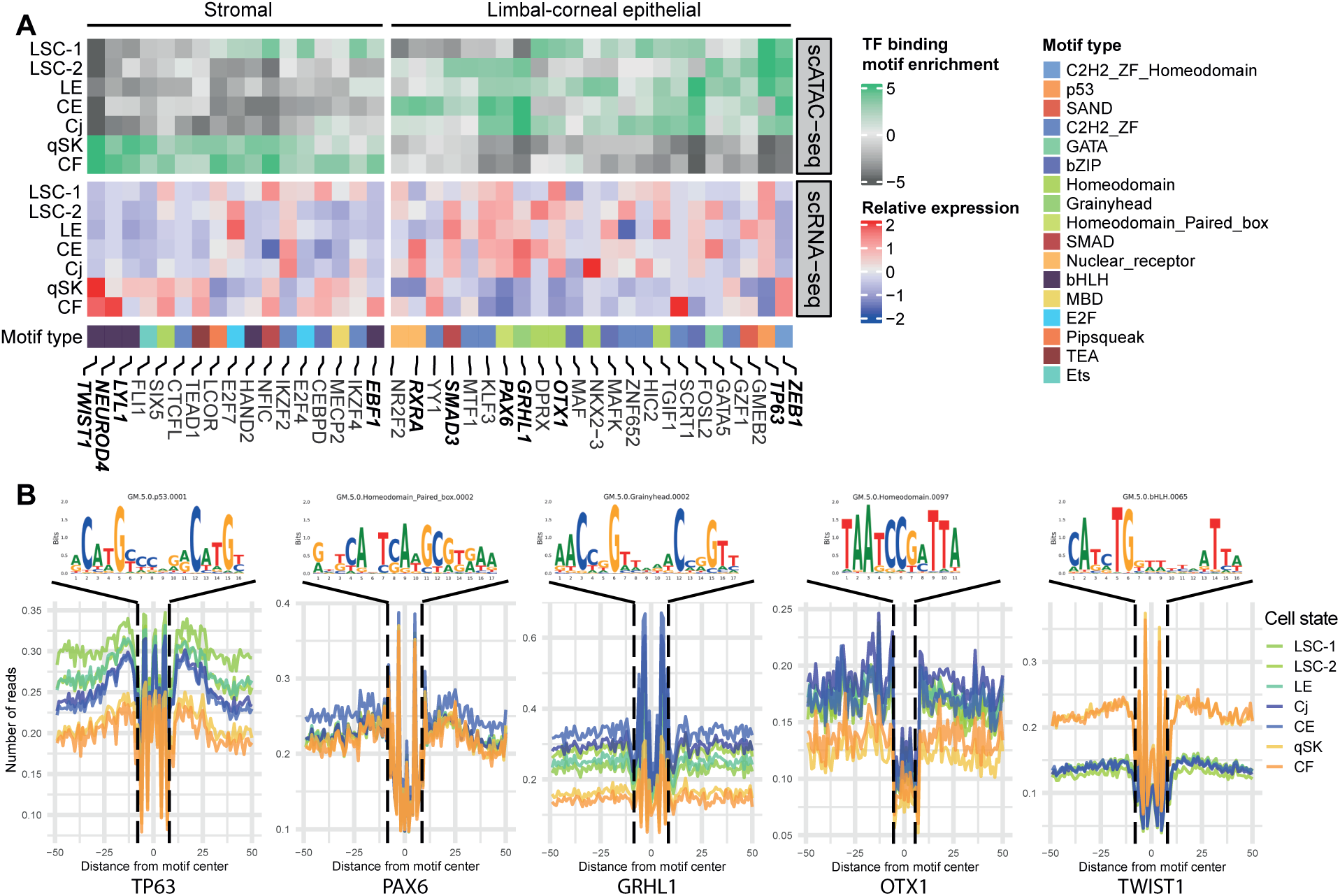
Motif enrichment and prediction of transcription factor binding in cell states of the human cornea: **A)** Heatmaps of the motif scores of the top 10 enriched TFs (upper panel) and of TF expression levels (lower panel) for each cell state. The bottom panel shows the type of associated motif. **B**) Examples of the consensus motif and transcription factor footprints from 5A. Names of cell states: LSC-1 = Limbal stem cells 1; LSC-2 = Limbal stem cells 2; LE = Limbal epithelium; CE = Central epithelium; Cj = Conjunctiva; qSK = Quiescent stromal keratocytes; CF = Corneal fibroblasts.

The group of TFs that had high TF binding enrichment scores in cell states of the non-epithelial branch included TWIST1, NEUROD4, LYL1, and EBF1 (Figure 5A), all of which are TFs associated with bHLH motifs. As expected, the region surrounding the TWIST1-associated motif consistently displayed high scATAC-seq signals in both qSK and CF (Figure 5E).

### Prediction of key transcription factors using gene regulatory network analysis

Using motif analysis on accessible chromatin regions predicts TFs that regulate gene expression in specific cell states, but it does not indicate the importance of TFs in driving cell state determination. To predict the importance of TFs driving corneal cell state identity, we leveraged our previously developed single-cell gene regulatory network (GRN) method, scANANSE^27^, which ranks the importance of a TF represented by an influence score in a specific cell state, as compared to another. For this comparison, embryonic stem cells^83^ were used against all cell states in the human corneal cell state meta-atlas (Figure 6A), a strategy that was previously shown to be effective for TF identification in similar cell states when applying the ANANSE pipeline^84^. In this analysis, we identified TFs with high influence scores shared in all cells (Figure 6B) but also those distinct to each branch (Figure 6C, 6D).

**Figure 6:**
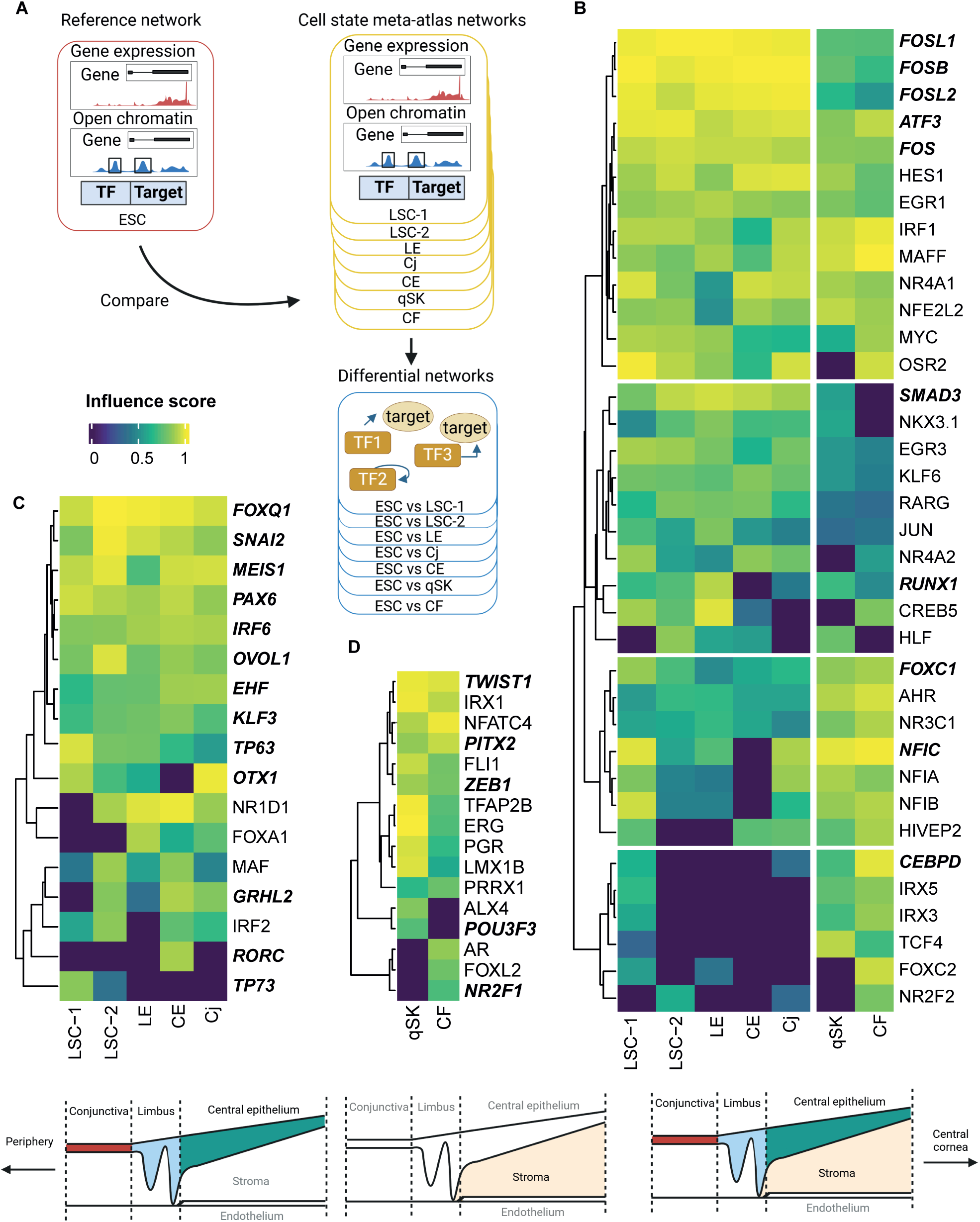
Prediction of key transcription factors controlling corneal cell states using gene regulatory network analysis of scANANSE. **A)** An in vitro single-cell hESC network was used as a reference against networks of all individual cell states in the corneal cell state meta-atlas. Black boxes enclosing peaks in open chromatin depict putative enhancers where TFs can bind. **B)** Heatmap of influence scores (> 0.8) of TFs shared between limbal/corneal epithelial and stromal cell states, predicted by scANANSE. **C)** Heatmap of influence scores (> 0.8) of TFs specific for limbal/corneal epithelial cell states, predicted by scANANSE. **D)** Heatmap of influence scores of the top 40 TFs specific for stromal cell states, predicted by scANANSE. Schematic pictures highlight the corresponding regions and cells in the cornea where TFs (B-D) play key roles. Names of cell states: LSC-1 = Limbal stem cells 1; LSC-2 = Limbal stem cells 2; LE = Limbal epithelium; CE = Central epithelium; Cj = Conjunctiva; qSK = Quiescent stromal keratocytes; CF = Corneal fibroblasts.

Among the shared TFs, we identified four groups (Fig 6B). The first group included FOSB, FOSL1, FOSL2, FOS and ATF3, TFs associated with the AP-1 complex^85^. They had high influence scores in both epithelial and non-epithelial branches, with the scores in the epithelial branch slightly higher than those in the non-epithelial branch. Among these five TFs important for regulating cell states in the human cornea, only FOSL2 was detected in our motif analysis (Figure 5A). The second group contained TFs with high influence scores in epithelial cell states and lower, but detectable scores in qSK and CF. This group contained SMAD3 and RUNX1, with known a function associated with limbal stem cells^24^. The third group contained TFs such as FOXC1^24^ and NFIC that had high influence scores in both branches but with higher scores in the non-epithelial branch; one of these, FOXC1 was not identified using motif enrichment analysis. The fourth group of TFs had high influence scores in cell states not specific to a branch. Remarkably, this group contained CEBPD that had high influence scores in LSC-1 as well as in qSK and CF, even though this TF was known to be associated specifically to limbal stem cells^86^.

Many identified TFs with high influence scores specific for limbal/corneal epithelial cell states are well-known epithelial TFs. These included EHF, KLF3, TP63 and PAX6 (Figure 6B), which are known for their role in corneal differentiation^21,87,88^, and FOXQ1, SNAI2, MEIS1 and OVOL1, which are known to be involved in WNT signaling^89–92^. Other TFs with high influence scores were OTX1, GRHL2, RORC and TP73. OTX1 exhibited highest influence scores in LSC-1 and Cj, which was consistent with our motif analysis (Figure 5A). GRHL2 showed a high influence score for LSC-2, CE and Cj. RORC showed a unique high influence score in CE, while TP73 had a high influence score in LSC-1 and a lower influence score in LSC-2.

Many of the identified non-epithelial specific TFs are less known in the cornea. Most of them, including PITX2, LMX1B, TWIST1, FLI1, ERG and ZEB1 exhibited high influence scores in both qSK and CF (Figure 6D), with TWIST1, FLI1, ERG and ZEB1 being consistent with motif analysis (Figure 5A). ALX4 and POU3F3 both had high influence scores only in qSK (Figure 6D), in line with their roles associated with maintaining mesenchymal identity^93,94^. To note, POU3F3 is one of the novel markers identified in this study, and its expression at the protein level was validated (Figure 2B and 3E). NR2F1 displayed a high score only in CF (Figure 6D), consistent with it being a fibroblast associated factor^95^.

Taken together, our analysis identified well-known and novel key TFs for corneal cell state determination. We also showed that corneal cell states are mostly driven by combinations of TFs, whereas a small number of TFs were cell state specific. scANANSE’s influence scores of TFs and general gene expression across these cell states can be interactively explored in a dashboard.

## Discussion

Understanding the precise cell states is pivotal for both *in vivo* and *in vitro* studies on development, pathogenesis and regeneration of the cornea. In this work, by integrating four scRNA-seq datasets, we annotated corneal cell states including previously unknown rare cell states, identified novel marker genes and created a corneal cell state meta-atlas. We demonstrated that the machine learning-based prediction pipeline cPredictor that applies the cell state meta-atlas as the reference can define cell states in various types of corneal scRNA-seq data. Furthermore, we characterized distinct combinations of key TFs controlling cell states of the human cornea, by integrating scATAC-seq with our scRNA-seq-based cell state meta-atlas. Both marker gene expression and TF influence scores can be interactively visualized through a web portal. This portal together with cPredictor will evolve continuously as an expanding resource for investigating human corneal cell states.

The corneal cell states characterized in this work are mostly consistent with known corneal cell states in literature^8,77,96,97^. Two populations of limbal stem cells were detected in our study; LSC-1 expressed the quiescence-related gene product *GPHA2*, whilst LSC-2 showed significant enrichment for genes linked to epidermal differentiation, indicating a more differentiated state of LSC-2 compared to LSC-1. These findings are in line with previous work showing one limbal stem cell population exhibiting quiescent stem cell traits and another active in corneal regeneration in mice^97^. Interestingly, our integrated corneal cell state meta-atlas supported the existence of a tiny number (<0.15%) of non-myelinating corneal Schwann cells (nm-cSC), which has been reported across various species^77,96,98^. Although the mechanisms behind nm-cSC function are not fully clear, these cells potentially play important roles in corneal wound healing^99^ and in cornea-associated diseases such as Familial Dysautonomia^100^ and neuropathic pain^101^. It is worth noting that the nm-cSC cell state was not identified in any of the individual scRNA-seq studies, likely due to its small cell number in the cornea.

We uncovered novel marker genes linked to corneal cell state identities in the integrated scRNA-seq cell state meta-atlas. Of note, we detected *SLC6A6* as a novel marker specific for the limbal stem cell state LSC-1, with high protein abundance limited to the corneal limbus. The presence of *SLC6A6* in the limbus is in line with a recent single-nucleus RNA-seq study^102^. *SLC6A6* plays roles in reducing reactive oxygen species (ROS) and in regulating the Wnt/β-catenin signaling pathway^103^, a signaling pathway known to be important in the limbus and in corneal development and differentiation^104,105^. In LSC-1, *SLC6A6* might be an important player in the Wnt/β-catenin pathway. In this work, several WNT-associated TFs FOXQ1, SNAI2, MEIS1 and OVOL1 were identified in our GRN analysis and showed high influence scores in LSC-1. Further investigation of this Wnt/β-catenin axis is warranted, as none of the WNT-associated TFs have been previously implicated in the regulation of limbal stem cells. Furthermore, *TNNC1*, a well-known cardiac cytoskeletal troponin gene^106^, was identified as a distinct marker gene for the corneal endothelium, and validated in our immunohistochemical staining of human corneas. Its potential role as a calcium sensor^106^ possibly needed for endothelial cell function^107^ in the human cornea needs to be further investigated.

The strength of the corneal cell state meta-atlas and its derived SVM-based machine learning pipeline cPredictor in the identification of rare cell states was demonstrated in our study. So far, only one scRNA-seq study of the human cornea^19^ was available for us to test its performance. cPredictor was able to predict 15 cell states with medium to high confidence scores, whereas only 12 cell states were predicted in the original study^19^. Furthermore, our corneal meta-atlas seems to be more robust as a corneal reference, as it was integrated from multiple datasets derived from different methods of cell retrieval^20^, and therefore contained more cell states than each individual study. We showed that the data from Collin and colleagues^14^ contributed to the majority of epithelial cells in the cell state meta-atlas. Additionally, this was the only study to retrieve mural cells, previously annotated as Fibroblast Corneal Endothelial Cells, probably due to tissue handling as they used bulk enzymatic sample disaggregation. In the study of Català and colleagues^13^, a dissection protocol that gently separates the cornea in multiple parts before disaggregation was carried out to retrieve high-quality corneal stromal cells, including stromal keratocytes.

Therefore, a small number of (transitioning) stromal keratocytes were only retrieved in this study^13^. How and why these biases exactly arise needs to be investigated in future studies. Nevertheless, our constructed corneal cell state meta-atlas together with cPredictor should be more robust to predict corneal cell states from different retrieval methods.

With its automated capabilities, cPredictor is easy to use. Our containerized software and command-line based approach enable ease-of-use for predicting adult corneal cell states in external datasets in a reliable prediction of adult corneal cell states. It is also straightforward to incorporate new datasets in cPredictor, if new scRNA-seq datasets of the human cornea are available. This may further strengthen its capability to detect rare cell states and marker genes. Additionally, cPredictor could potentially incorporate other meta-atlases as training datasets, e.g. from mouse cornea data or other tissues, expanding its applicability beyond human cornea research.

Furthermore, applying cPredictor allowed us to investigate cell states in iPSC-derived corneal organoids, using the corneal cell state meta-atlas as the reference. As the meta-atlas was generated from four scRNA-seq datasets derived from adult human corneas^13–15,17^, cell states in organoids were compared to those in human adult corneas. Our results confirmed that current iPSC-derived corneal organoids do not yet resemble the cell states in adult corneas. Our predictions showed that conjunctival (Cj) and corneal fibroblast (CF) cell states in 4-month corneal organoids were most similar to the adult cell states, represented by certainty scores and detected marker genes. This suggests that studying Cj and CF using organoids could give relevant information on their cell functions in the cornea. However, many cell states, like endothelial cells, still show large differences, as compared to those in the adult cornea. The clear differences in the cell states between the adult human corneas and iPSC-derived corneal organoids suggest that corneal organoid generation needs further improvement, to be able to fully mimic the function of the cornea, potentially via maturation. That maturation may be one of the bottlenecks is supported by the observations that organoids generated at the end of month 1-3 were less similar to the reference, as compared to the 4-month-old organoids.

Prolonging the culturing time of the organoids could one strategy. Alternatively, proper environmental cues, such as signaling molecules or mechanical force, could improve the maturity or functionality of cells in corneal organoids. Such information may be derived from further in-depth studies on the similarities and differences between the human adult corneas and iPSC-derived corneal organoids.

The identified TFs through scANANSE GRN analysis revealed many well-known key (corneal) epithelial TFs, including PAX6^108^, TP63^22^, SMAD3^24^, GRHL2^109^ and FOSL2^84^ in all limbal/corneal epithelial cell states. RXRA exhibits a high TF motif binding enrichment score in CE, and RORC, a downstream target of RXRA^110^, had a high influence score exclusively in CE. ATRA, known for dimerizing with RXRA, promotes corneal epithelial cell integrity and transparency^111^. TFs in non-epithelial cell states qSK and CF included PITX2, POU3F3, ALX4, LMX1B, TWIST1, FLI1 and ERG. Except PITX2 that is known to play important roles in neural crest specification^112^ and proper development of the corneal stroma^113^, other TFs are less studied in the cornea. POU3F3, of which the protein was localized in the cell nuclei of the corneal stroma, was identified as a novel TF and marker gene for qSK in our study. However, protein expression of POU3F3 was also detected in the cytoplasm in the corneal epithelium, where POU3F3 is probably not functioning as a TF. LMX1B is known for its function in periocular mesenchyme-derived cell identity^114,115^, similar to PITX2, but unknown in the cornea. In addition, the roles of TWIST1, FLI1 and ERG, in corneal stromal cell states are completely unexplored. Further research on the function of these TFs is warranted.

In summary, we show that the corneal cell state meta-atlas can serve as a reliable reference for annotating and predicting corneal cell states, e.g., in dissecting the difference between healthy and diseased cells. The easy-to-use computational pipeline cPredictor and the identified marker genes and key TF in various cell states of the human cornea provide a rich resource for follow-up research on corneal biology and regeneration.

## Acknowledgements

The authors thank ETB-BISLIFE: Multi Tissue Center for providing research-grade human corneas. The cartoons in Figure 3A, Figure 6A–D and Supplemental Figure 4A were created with BioRender.com.

The authors also wish to thank the following funding sources for their financial support to this study: ZonMw Open (09120012010039, to H.Z., J.A.A., D.L.C., M.D., V.L.S.L, S.F.), European Joint Programme on Rare diseases (EJPRD20-135, to H.Z., D.L.C.), and funding from Velux Stiftung (to H.Z.).

## Author contributions

Conceptualization: H.Z., J.A.A., and D.L.C.; Tissue sectioning: M.S.F. and S.F., IF: S.F.; scRNA-seq data analysis: J.A.A.; machine learning model conceptualization: J.A.Y.R. and J.A.A.; Construction of the cPredictor machine learning pipeline: J.A.A.; writing – original draft: J.A.A.; writing – review and editing: J.A.A., S.F., M.S.F., J.G.A.S., D.L.C., J.A.Y.R., M.D., V.L.S.L., R.Y. and H.Z.

## Methods

### Ethical statement

In this study, two human donor corneas deemed unsuitable for transplantation were used for immunofluorescence analysis. The corneal tissues were obtained from two male donors, aged 70 and 71, from the ETB-BISLIFE Multi-Tissue Center (Beverwijk, the Netherlands). These tissues were preserved in Organ culture media at 31°C. The composition of the media included the following: minimum essential medium (MEM) supplemented with 20 mM HEPES, 26 mM sodium bicarbonate, 2% (v/v) newborn calf serum (Thermo Fisher Scientific), 10 IU/mL penicillin, 0.1 mg/mL streptomycin, and 0.25 μg/mL amphotericin. Both donor tissues had no history of ocular disease or infection, including HIV or hepatitis B.

### Immunofluorescence

Corneas were halved transversely, embedded in paraffin, and fixed in 4% paraformaldehyde at room temperature (RT) for 10 minutes. To preserve tissue morphology, 10 μm thick sections were cut consecutively on adhesive cryofilm type 3C (16UF) using a modified Kawamoto method with slight adjustments^116^. For antigen retrieval, tissue sections were incubated in boiling citrate buffer with 0.05% Tween 20 at 95°C (pH 6) for 20 minutes or treated with Pepsin C (Cat. No. ab64201; Abcam) for 10 minutes at RT. Sections were then permeabilized with PBS containing 0.2% Triton X-100 (Cat. No. T8787; Sigma-Aldrich) for 10 minutes at RT. Blocking was performed in PBS containing 10% goat serum and 2% bovine serum albumin (PBS-2% BSA) for 1.5 hours at RT, followed by incubation with primary antibodies (Supplemental Table 1) diluted in PBS-2% BSA overnight at 4°C in a humidified chamber. The following day, sections were washed three times with PBS-T and incubated with fluorescently conjugated secondary antibodies (goat anti-rabbit A488 and goat anti-mouse A647, both at 1:1000 dilution; Thermo Fisher Scientific) for 1 hour at RT in the dark. Sections were then washed again three times with PBS-T and counterstained with DAPI (1:2000) for 10 minutes at RT. After a final set of three washes, coverslips were mounted using Fluoromount G (Thermo Fisher Scientific). Imaging was performed on an automated inverted Nikon Ti-E microscope equipped with a Lumencor Spectra light source, an Andor Zyla 5.5 sCMOS camera, and an MCL NANO Z200-N TI z-stage, using a CFI S Plan Fluor ELWD 40X objective. Image analysis was carried out using NIS-Elements software.

### Data and code availability

Datasets used in this study are from previous publications and currently publicly available under the Gene Expression Omnibus (GEO) identifiers GSE155683, GSE186433, GSE153515, GSE147979, GSE178379, GSE218123 and GSE240458^13–15,17,19,81,83^ were used. All these datasets contain single-cell RNA-seq data. Among these datasets, GSE155683 and GSE178379 also contain single-cell ATAC-seq data. The full processing workflow and documentation of code used for Python and R are available in the GitHub repository: https://github.com/Arts-of-coding/Cell-States-and-Key-Transcription-Factors-of-the-Human-Cornea-through-Integrated-Single-Cell-Omics. GRCh38 downloaded with Genomepy version 0.10.0 ^115^, was used for all downstream analyses. An interactive web app was developed to visualize corneal scRNA-seq meta-atlas, gene expression, and influence scores of TFs. The interactive web app is available at: https://huggingface.co/spaces/Arts-of-coding/corneal_cell_state_meta_atlas. Single-cell objects of the meta-atlas and files used by cPredictor in this study can be downloaded from Zenodo^117^: https://doi.org/10.5281/zenodo.7970736. Exploration of pseudobulk scATAC-seq is available using the link below in the UCSC genome browser track hub: https://mbdata.science.ru.nl/jarts/scATAC_Corneal_meta-atlas/ATAC-seq_trackhub.hub.txt.

### Pre-processing single-cell RNA-seq and single-cell ATAC-seq

Datasets were downloaded from GEO with Seq2Science^118^. The 10X .sra files were split into fastq files with Seq2Science. Next, Snakemake pipelines were developed to pre-process multiple datasets with cellranger. Using these pipelines, Cellranger count was run with Cellranger 7.0.1^119^, with default parameters and with hg38 to retrieve the matrix, barcodes and features files necessary for downstream analysis. For GSE178379 specifically, Cellranger-arc 2.0.2 was run to retrieve these files for multi-omics data. Comparable to scRNA-seq pre-processing, a Snakemake pipeline containing Cellranger-atac count was run with Cellranger-atac 2.0.0^120^ to generate both the position sorted bam file and the singlecell.csv file, containing barcode information, necessary for snapATAC^121^ analysis in Bash and R.

### Quality control of scRNA-seq and scATAC-seq

scRNA-seq datasets were analyzed in R with Seurat version 4.0.2^122^, as described in the GitHub repository. scRNA-seq cells were selected with a minimum count of 2000, a feature number higher than 1000 and a mitochondrial percentage lower than 30 percent. Expected doublets were removed with DoubletFinder version 2.0.3^29^. scATAC-seq datasets were processed in R with SnapATAC version 2.0^116^. Due to the higher sparsity of the data compared to scRNA-seq, a strict quality control was used. Only cells with a mitochondrial ratio lower than 30 percent were selected. Additionally, the fragment threshold number for each cell was set between 5000 and 500.000. To minimize the number of doublets that could skew the data, a maximum of 500.000 for the fragment and a number between 3.75 and 5 for the unique count were chosen. A Log10 coverage above 3.6 and a promoter ratio between 2 and 8 were used and cells with a duplicate ratio higher than 0.8 and a low mapping quality value higher than 5000 were excluded. scRNA-seq data from GSE218123 and GSE240458 were analyzed in Python with Scanpy version 1.9.2, as described in https://github.com/Arts-of-coding/Cell-States-and-Key-Transcription-Factors-of-the-Human-Cornea-through-Integrated-Single-Cell-Omics. A minimum count of 900 and a mitochondrial percentage lower than 20 percent were used as described by the study of Swarup^81^.

### Integration and clustering of scRNA-seq datasets

Clustering in each dataset was conducted with specific parameters from dimension numbers (Elbow plot) and with clustering resolution using Clustree version 0.4.4^123^. Clustering on scRNA-seq data from the studies of Català, Collin, Gautam and Li was performed with Leiden clustering^32^, using 30, 16, 18 and 24 dimensions and a clustering resolution of 0.15, 0.1, 0.15 and 0.1 respectively. The four datasets were integrated with scVI (scvi-tools version 0.14.0)^30^. The parameters for the Variational Autoencoder were selected as 2 for “n_layers”, 30 for “n_latent” and “nb” for “gene_likelihood”. Leiden clustering was performed on the integrated data using a clustering resolution of 0.3.

### GO-term enrichment analysis

We performed gene ontology enrichment on all marker genes found in each cell state by use of the org.Hs.eg database^124^ for biological processes, with the cut-off of BH-adjusted p-values <0.05^125^. Similarly, GO-terms that had a similarity score regarding overlapping genes of 0.5 (range 0-1) for overlapping genes were joined together using the function simplify from ClusterProfiles version 3.0.4. Further selection on the GO-terms was performed by selecting terms with a minimum gene ratio of 0.1 and terms consisting of at least 200 genes per term. The top 3 highest counted GO terms were visualized.

### Pathway enrichment analysis

Pathway enrichment was performed with PROGENy (decoupler-py version 1.6.0), a tool that showed enrichment for 14 curated signal-transduction pathways^36^. The top 500 genes for each pathway were selected and no permutations were added. Activity for these curated pathways was inferred for each single cell by running a multivariate linear model from decoupleR. Subsequent visualization was performed in Python.

### cPredictor classifier training and prediction with the corneal meta-atlas

All scripts for iterative feature (gene) selection, hyperparameter optimization and model explainability can be found under https://github.com/Arts-of-coding/Cell-States-and-Key-Transcription-Factors-of-the-Human-Cornea-through-Integrated-Single-Cell-Omics The training dataset supplied to cPredictor^126^, using support vector machines from Scikit-learn (Sklearn)^82^, was the corneal cell state meta-atlas. Raw scRNA-seq count data was transformed with “log1p” from numpy and MinMax scaling from Sklearn. A 5-fold cross-validation was run on the corneal meta-atlas dataset to estimate generalized model performance. Within the automated pipeline, metrics were logged and calibration for each cell state was generated in a one-versus-all (single-class) fashion. Features (genes) were selected through an iterative approach of gene elimination, starting with expressed genes in the corneal meta-atlas (Supplemental Figure 4A). In each round the top-n highly variable genes and the top-n explainable genes were taken along if the model showed good calibration for that specific cell state. In the end, top explainable genes *MALAT1*, *XIST*, *SRY*, *MT1Z*, *GAS5*, *BTG2*, *CD9* and *SNHG32* were removed as they contributed to model explanations being either expressed in all cell states or being sex-chromosome specific which negatively affects model predictions. After feature selection, model hyperparameter optimization was performed using Monte-Carlo simulations^127^. The hyperparameters in the final model were chosen based both on the optimal ROC-AUC and accuracy scores across classes and can be found under: https://huggingface.co/Arts-of-coding/meta-atlas-cornea-SVM. The Docker container “artsofcoding/cpredictor:v0.4.5_cpredictor_hcornea_v1” was used for cell state predictions on the data from Maiti^19^ and Swarup^81^.

### Prediction of scATAC-seq cells from scRNA-seq data

After annotating the cell clusters in scRNA-seq and performing quality control as described, scATAC-seq cell clusters were imputed with Seurat label transfer with a minimum prediction score of 0.4. The position-sorted bam and singlecell.csv files were used to generate snap files, as described in the SnapATAC package^121^.

### Bam file generation, transcription factor footprinting and motif analysis

Cell barcodes were retrieved in R to generate a pseudobulk profile of ATAC-signal for each cell state. For downstream analysis, we performed splitting of full bam files by cell states in three of the four datasets, which showed high quality. We performed this barcode splitting for the imputed cell populations consisting of more than 100 cells in scATAC-seq. Bam files were split for cell populations with bamslice based on corresponding cell barcodes with cellranger-dna version 1.1.0 to enable peak calling. Peaks for motif analysis were generated from the .bam files with peak calling software MACS2^128^. Both the bam and Narrowpeak files were used to generate accessible summit files and peak count files with motif analysis software Gimmemotifs^129^. These two files were used in the q-quantile normalization of peak count files and were used for downstream scANANSE analysis^27^. The output of Gimmemotifs and Z-score expression of transcription factors were correlated in R to determine which transcription factor was most likely to bind a motif. Footprinting analysis was performed on bam files, bed files and Narrowpeak files of cell states with HINT-ATAC using RGT version 0.13.2^130^. Briefly, rgt-hint footprinting function was used on our Narrowpeak files. Next, the rgt-motif analysis matching function was run on our created bed files. The motifs were matched against the full Gimme Motifs database. Lastly, the footprint files were generated using the rgt-hint differential function. The output files were imported in R to trim the distance from the center of the motif to 50bp upstream as well as 50bp downstream.

### Gene regulatory network analysis of single-cell states

scANANSE was performed on single-cell clusters in a pseudo-bulk fashion by using the package AnanseScanpy^27^. In short, GRNs of single-cell populations consisting of more than 100 cells present in scATAC-seq were generated. The hg38 genome and the combined pseudo bulk peak matrix, generated with the function export_ATAC_scANANSE of all populations were used for performing ANANSE binding. The output network file from ANANSE binding was used, next to the hg38 as the reference genome for the ANANSE network step. Due to the nature of the 3’ sequencing, CPM values were used as input for generating the network files, generated with the function export_RNA_scANANSE from the single-cell object.

Differential gene expression across single cells was performed by using the export_DEGs_scANANSE function. For ANANSE influence scores, two different networks were compared. A network composed of scATAC-seq and scRNA-seq from human naïve embryonic stem cells^83^ (see data availability) was used as a reference for each of our cell states in the corneal meta-atlas. All corneal cell state meta-atlas networks were compared to this network in the scANANSE influence step. These binding, network and influence steps were performed with ANANSNAKE, as described in the scANANSE workflow.

### Declaration of the use of AI in the writing process

During preparation of this work, generative AI models were used to proofread and improve the fluency of the manuscript text. All text was carefully reviewed and edited as needed by the authors, who take full responsibility of all contents of this publication.

