## Supplemental Information for "Prediction of Cell States and Key Transcription Factors of the Human Cornea through Integrated Single-Cell Omics Analyses"

### 1. Supplemental figures and tables

#### 1.1. Figures S1-S6

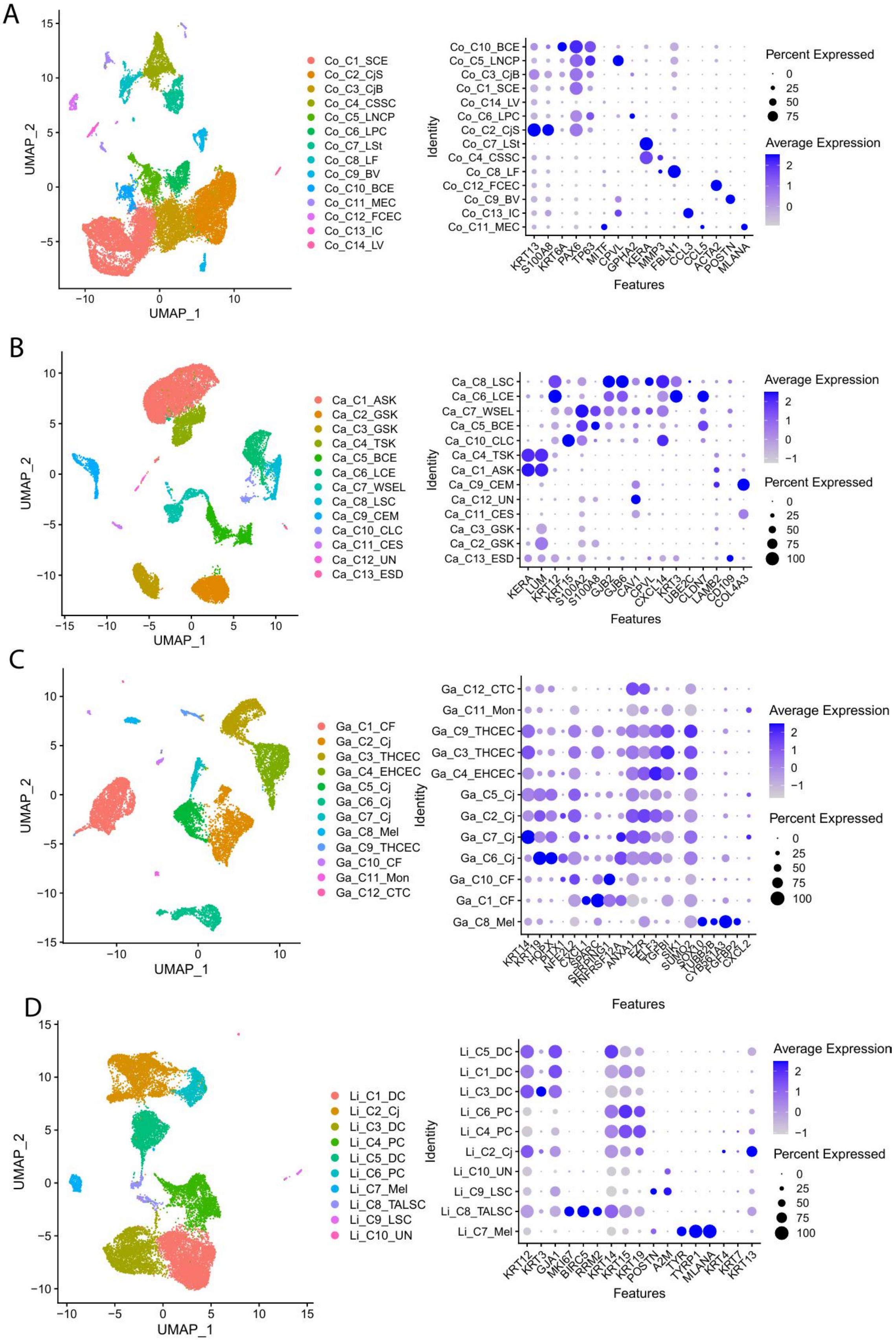

***Supplemental Figure 1: Defining cell states in the human cornea across individual corneal studies based on proposed marker genes****. UMAPs of defined cell states and dot plots of marker genes in the study of (A) Collin (B) Català (C) Gautam (D) Li. The original identified cell states in individual studies are listed below.*

*Names of cell states (A): BCE= Basal corneal epithelium; LNCP = Limbal neural crest-derived progenitor cells; CjB= Conjunctival basal; SCE = Superficial corneal epithelium; LV = Lymphatic vessels; LPC = Limbal progenitor cells; CjS = Conjunctival superficial; LSt = Limbal stromal cells; CSSC = Corneal stromal stem cells; LF = Limbal fibroblasts; FCEC = Fibroblast corneal endothelial cells; BV = Blood vessels; IC = Immune cells; MEC = Melanocytes and endothelial cells.*

*Names of cell states (B): LSC = Limbal stem cells; LCE = Limbal corneal epithelium; WSEL = Wing superficial epithelial limbal cells; BCE= Basal corneal epithelium; CLC = Corneal limbal cells; TSK = Transitioning stromal keratocytes; ASK = activated stromal keratocytes; CEM= Corneal endothelium migratory; UN = Uncharacterizable; CES = Corneal endothelium stationary; GSK = General stroma keratocytes; ESD = Epidermal-stromal doublets.*

*Names of cell states (C): CTC = Cytotoxic T-cells; Mon = Monocytes; THCEC = TGFβ-high corneal epithelial cells; EHCEC = ELF3-high corneal epithelial cells; Cj= Conjunctiva; CF = Corneal fibroblasts; Mel = Melanocytes.*

*Names of cell states (D): DC = Differentiated cells; PC = Progenitor cells; Cj= Conjunctiva; UN = Uncharacterizable; LSC = Limbal stem cells; TALSC = Transient amplifying limbal stem cells; Mel = Melanocytes.*

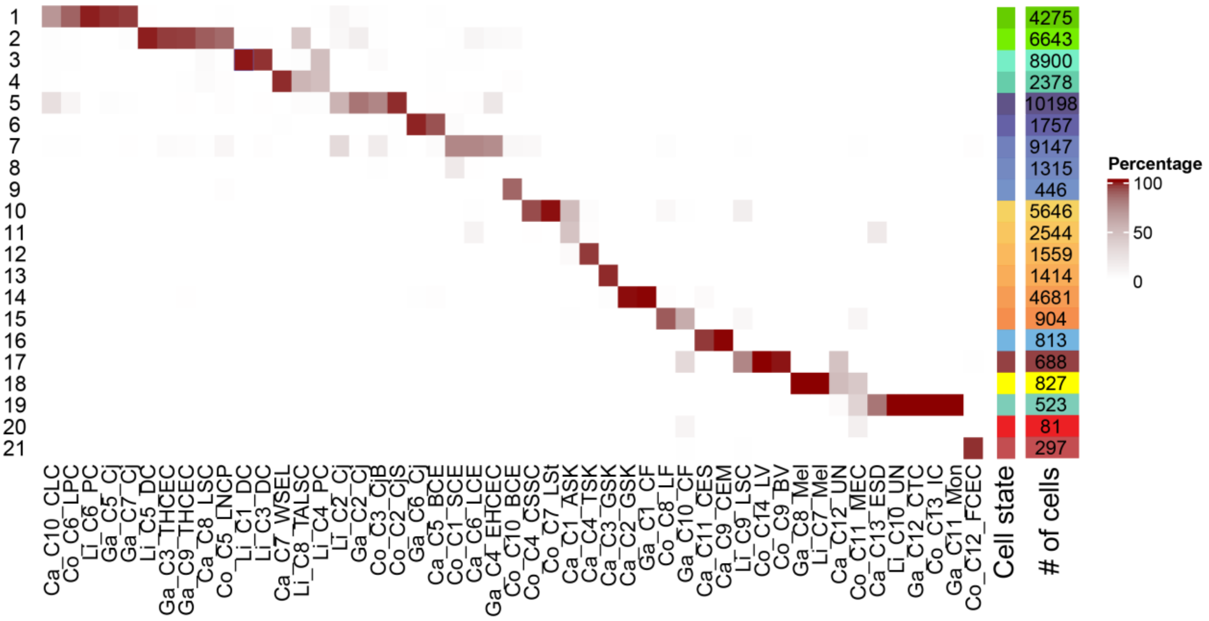


***Supplemental Figure 2: Contingency table of integrated cell clusters with defined cell states across studies****. Cell numbers and cell states in the integrated data are annotated on the right, and the original cell states in individual studies are indicated on the x-axis. Ca stands for Català, Co stands for Collin, Ga depicts cells from Gautam and Li indicates cells from Li. C indicates the cluster number across individual studies. The cell origins from A are depicted on the right, together with the total number of cells.*

*Names of cell states from the individual studies: CLC = Corneal limbal cells; LPC = Limbal progenitor cells; PC = Progenitor cells; Cj= Conjunctiva; DC = Differentiated cells; THCEC = TGFβ-high corneal epithelial cells; LSC = Limbal stem cells; LNCP = Limbal neural crest derived progenitor cells; WSEL = Wing superficial epithelial limbal cells; TALSC = Transient amplifying limbal stem cells; PC = progenitor cells; CjB= Conjunctival basal; CjS = Conjunctival superficial; BCE= Basal corneal epithelium; SCE = Superficial corneal epithelium; EHCEC = ELF3-high corneal epithelial cells; CSSC = Corneal stromal stem cells; LSt = Limbal stromal cells; ASK = activated stromal keratocytes; TSK = Transitioning stromal keratocytes; GSK = General stroma keratocytes; CF = Corneal fibroblasts; LF = Limbal fibroblasts; CES = Corneal endothelium stationary; Corneal endothelium migratory; LV = Lymphatic vessels; BV = Blood vessels; Mel = Melanocytes; UN = Uncharacterizable; MEC = Melanocytes and endothelial cells; ESD = Epidermal-stromal doublets; CTC = Cytotoxic T-cells; IC = Immune cells; Mon = Monocytes; FCEC = Fibroblast corneal endothelial cells.*

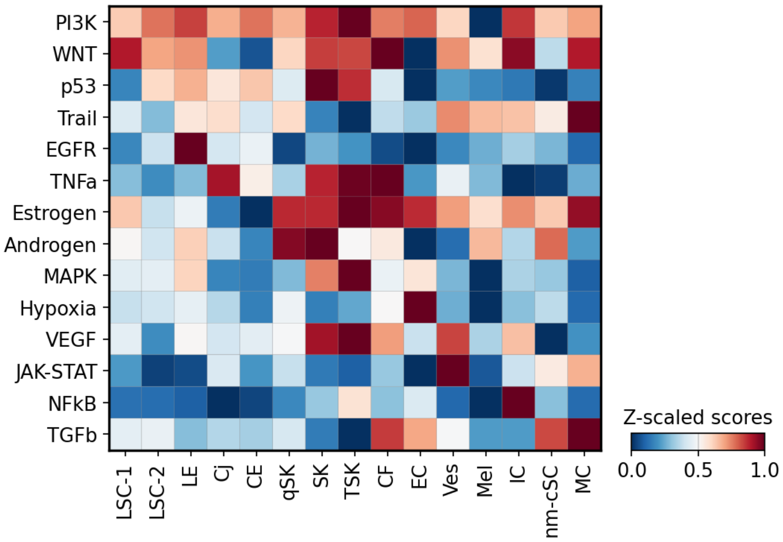


***Supplemental Figure 3: Pathway analysis of cell states in the corneal meta-atlas with PROGENy.** PROGENy scores scaled between 0 and 1 for 14 curated pathways based on the expression of genes involved in the respective pathways.*

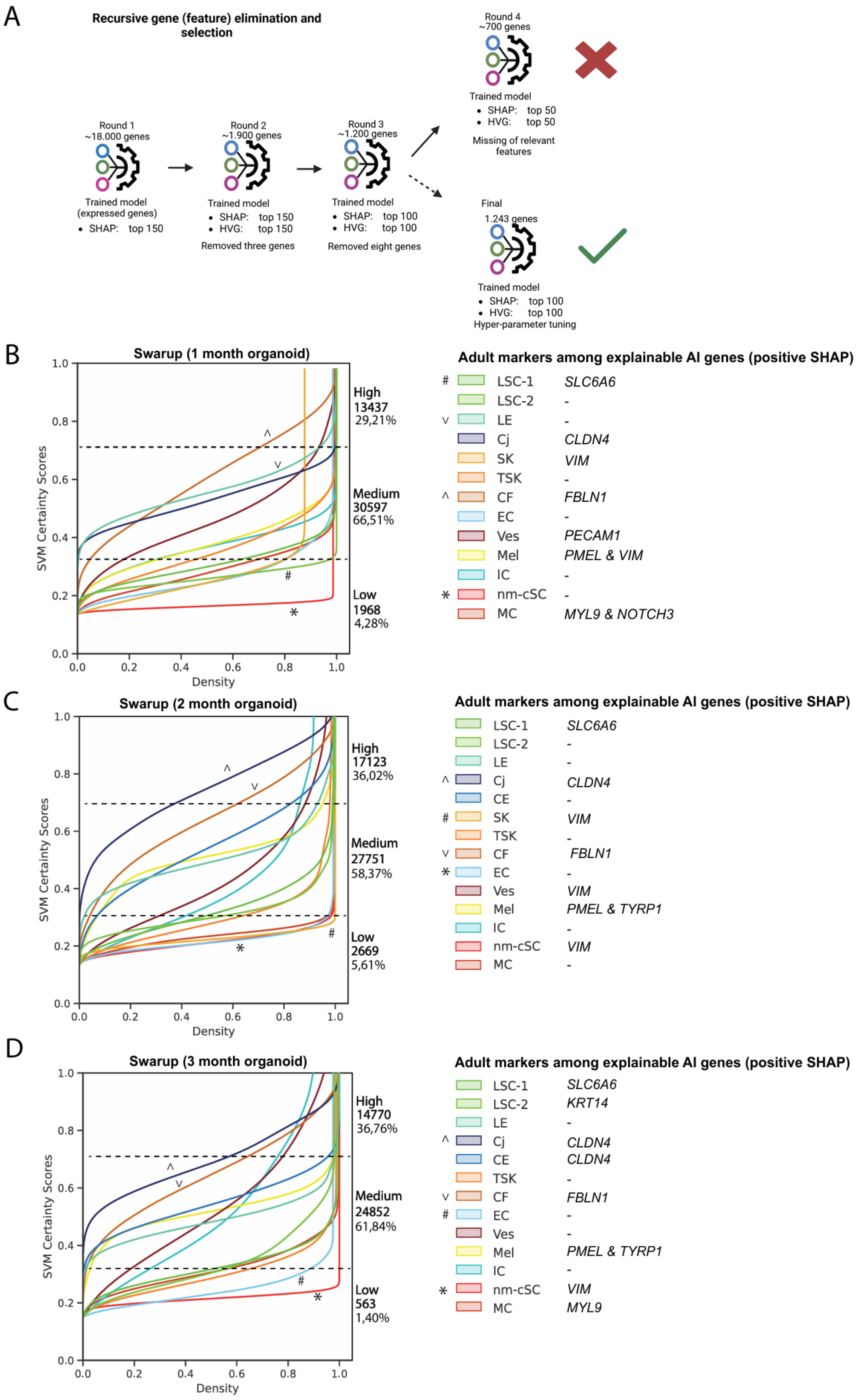
A
Recursive gene (feature) elimination and selection
Round 1
~18.000 genes
Trained model (expressed genes)
• SHAP: top 150
Round 2
~1.900 genes
Trained model
• SHAP: top 150
• HVG: top 150
Removed three genes
Round 3
~1.200 genes
Trained model
• SHAP: top 100
• HVG: top 100
Removed eight genes
Round 4
~700 genes
Trained model
• SHAP: top 50
• HVG: top 50
Missing of relevant features
Final
1.243 genes
Trained model
• SHAP: top 100
• HVG: top 100
Hyper-parameter tuning
B
Swarup (1 month organoid)
SVM Certainty Scores
Density
High
13437
29,21%
Medium
30597
66,51%
Low
1968
4,28%
Adult markers among explainable AI genes (positive SHAP)
### LSC-1 SLC6A6
LSC-2 -
v LE -
Cj CLDN4
SK VIM
TSK -
^ CF FBLN1
EC -
Ves PECAM1
Mel PMEL & VIM
IC -
* nm-cSC -
MC MYL9 & NOTCH3
C
Swarup (2 month organoid)
SVM Certainty Scores
Density
High
17123
36,02%
Medium
27751
58,37%
Low
2669
5,61%
Adult markers among explainable AI genes (positive SHAP)
LSC-1 SLC6A6
LSC-2 -
LE -
^ Cj CLDN4
CE -
### SK VIM
TSK -
v CF FBLN1
* EC -
Ves VIM
Mel PMEL & TYRP1
IC -
nm-cSC VIM
MC -
D
Swarup (3 month organoid)
SVM Certainty Scores
Density
High
14770
36,76%
Medium
24852
61,84%
Low
563
1,40%
Adult markers among explainable AI genes (positive SHAP)
LSC-1 SLC6A6
LSC-2 KRT14
LE -
^ Cj CLDN4
CE CLDN4
TSK -
v CF FBLN1
# EC -
Ves -
Mel PMEL & TYRP1
IC -
* nm-cSC VIM
MC MYL9

***Supplemental Figure 4: Machine learning-based predictions of human corneal cell states in scRNA-seq datasets. A)*** *Schematic overview of recursive gene elimination. Each round model predictions were explained on their own training data and the top explainable AI (SHAP) genes were added into the pool of the top highly variable genes (HVG) for each cell state until the model performed well and still retained important marker genes for the cornea. In the final round the top 100 SHAP and HVGs were used, and hyper parameter tuning was performed.* ***B-D)*** *Prediction certainty plots of corneal cell states on 1-month-old (B), 2-month-old (C) and 3-month-old organoids from Swarup[80] (D). Left, the x-axis shows the cumulative kernel densities, and the y-axis depicts the model confidence (SVM certainty score). The numbers and percentages of cells corresponding to low (< 0.3), medium (>0.3 and <0.7) and high (> 0.7) certainty scores in each dataset are depicted next to the plots. ^/v indicate cell states most similar and */# indicate cell states least similar to cell states from the corneal meta-atlas. Right, corneal markers among the top ten explainable AI genes driving model's decisions for each of the adult corneal cell states determined by their SHAP positive values are shown.*

A

| | gene 1 | gene 2 | gene 3 | gene 4 | gene 5 | gene 6 | gene 7 | gene 8 | gene 9 | gene 10 |
|---|---|---|---|---|---|---|---|---|---|---|
| LSC-1 | MT1X | **KRT14** | **SLC6A6** | **S100A2** | DST | **KRT15** | PMAIP1 | MIR205HG | ID3 | **GPHA2** |
| LSC-2 | ALDH3A1 | IGFBP6 | **KRT14** | NQO1 | MIR205HG | TGFBI | HTRA1 | TKT | CAMK2N1 | COL17A1 |
| LE | RPS10 | FABP5 | TNFRSF12A | HMGA1 | **AREG** | SFN | SH3BGRL3 | **KRT14** | SOD2 | FOSL1 |
| Cj | **CLDN4** | AQP3 | PTGS2 | PERP | TGFBI | TACSTD2 | BTG1 | S100A11 | NEAT1 | CD55 |
| CE | **KRT3** | CD24 | ADIRF | PTGDS | **KRT12** | **CLDN4** | APOBEC3A | FABP5 | PERP | UPK3BL1 |
| qSK | ANGPTL7 | RPS4Y1 | **LUM** | **KERA** | HMOX1 | HTRA1 | DDX3Y | STEAP4 | PTGDS | CD81 |
| SK | DCN | SAA1 | ITGBL1 | CD81 | MRPS24 | FOSL1 | **VIM** | TNFRSF12A | **MMP3** | IFITM2 |
| TSK | RPS4Y1 | MT1X | PTX3 | MT2A | SSR3 | DDX3Y | SERPINA3 | DCN | LUCAT1 | MYDGF |
| CF | CXCL1 | NNMT | IGFBP5 | SAA1 | **FBLN1** | COL12A1 | HTRA1 | IGFBP7 | RND3 | IGFBP4 |
| EC | MGP | PTGDS | **SLC4A11** | **COL4A3** | ENO1 | SFRP1 | ITM2C | CCDC144A | APP | CA12 |
| Ves | CLDN5 | GNG11 | **VIM** | **ACKR1** | **PECAM1** | TGFBR2 | KLF2 | TCF4 | CCL14 | SPARCL1 |
| Mel | DCT | **PMEL** | **TYRP1** | **MLANA** | APOE | QPCT | **VIM** | CD63 | MITF | TRPM1 |
| IC | SRGN | **CCL4** | **CCL3** | CYBA | **VIM** | STK4 | CCL3L1 | BTG1 | DUSP2 | HLA-B |
| nm-cSC | **CDH19** | GPM6B | NRXN1 | **SCN7A** | LGI4 | KCNMB4 | RPS4Y1 | PLP1 | SAMHD1 | WNT6 |
| MC | TAGLN | ID4 | **MYL9** | NR2F2 | C11orf96 | RGS5 | **NOTCH3** | SPARCL1 | TPM1 | PPP1R12A |

B

| | gene 1 | gene 2 | gene 3 | gene 4 | gene 5 | gene 6 | gene 7 | gene 8 | gene 9 | gene 10 |
|---|---|---|---|---|---|---|---|---|---|---|
| LSC-1 | **S100A2** | MT1X | MIR205HG | KRT17 | DST | ID3 | MT-CO3 | **KRT15** | SNHG6 | SNHG29 |
| LSC-2 | IGFBP6 | **BCAM** | MIR205HG | STMN1 | ANXA1 | CAMK2N1 | TKT | KRT5 | IER3 | PLEC |
| LE | RPS10 | TNFRSF12A | GAPDH | SFN | SH3BGRL3 | FABP5 | KRT6A | HMGN3 | HMGA1 | SRSF5 |
| Cj | AQP3 | S100A11 | **KRT13** | **CLDN4** | BTG1 | NEAT1 | PERP | KRT17 | ITGA2 | DSC2 |
| CE | CD24 | KRT17 | S100A14 | KRT5 | ID1 | TKT | S100A9 | KRT13 | FTH1 | IGFBP2 |
| SK | **VIM** | ANXA5 | ANXA1 | EIF5A | TNFRSF12A | MRPS24 | SNHG7 | KLF6 | RPS4X | CEBPD |
| TSK | MT1G | MT-CYB | MT1X | MT-ND2 | SNHG29 | SSR3 | NUDC | CCDC85B | MTDH | CCNI |
| CF | TIMP1 | **VIM** | **FBLN1** | KRT17 | RND3 | FTL | SNHG5 | EIF1B | AKAP12 | CYP1B1 |
| EC | TTN | ENO1 | PTGDS | **TNNC1** | IER3 | NFKBIA | ID1 | MT-CO1 | APOE | APP |
| Ves | CLDN5 | **VIM** | GNG11 | CRIP2 | ADGRL4 | TCF4 | ARHGAP29 | EGFL7 | **PECAM1** | HLA-E |
| Mel | DCT | **PMEL** | APOE | **VIM** | **TYRP1** | CD63 | SDCBP | CD59 | KLF6 | STMN1 |
| IC | SRGN | **CCL4** | **CCL3** | PTPRC | LAPTM5 | STK4 | B2M | SAMSN1 | CYBA | CD44 |
| nm-cSC | **VIM** | CALM2 | WSB1 | CRYAB | GPM6B | HSP90AB1 | NRXN1 | CST3 | CNN3 | RHOB |
| MC | **MYL9** | A2M | TAGLN | TPM1 | CALD1 | PPP1R12A | **ACTA2** | SEPTIN7 | ID4 | DSTN |

C

| | gene 1 | gene 2 | gene 3 | gene 4 | gene 5 | gene 6 | gene 7 | gene 8 | gene 9 | gene 10 |
|---|---|---|---|---|---|---|---|---|---|---|
| LSC-1 | MT1X | **KRT15** | **SLC6A6** | **KRT14** | TXNIP | DST | **S100A2** | **GPHA2** | ID3 | ATF3 |
| LSC-2 | ALDH3A1 | **BCAM** | **KRT14** | STMN1 | IGFBP6 | FTH1 | NQO1 | TKT | ANXA1 | **KRT12** |
| LE | RPS10 | GAPDH | SFN | S100A10 | TNFRSF12A | HMGN3 | SH3BGRL3 | FABP5 | FOSL1 | CST3 |
| Cj | S100A11 | **KRT13** | **CLDN4** | AQP3 | KRT19 | CD55 | **KRT15** | MGST1 | BAG1 | FTH1 |
| CE | CD24 | S100A14 | KRT17 | ID1 | PTGDS | MT1X | MGARP | ELF3 | COX7B | SLC20A1 |
| SK | RPS4X | **VIM** | ANXA5 | EIF5A | IFITM2 | ANXA1 | HSPD1 | HSPB1 | S100A10 | TNFRSF12A |
| TSK | MT1G | MT1X | MT2A | NOP53 | MT1E | EEF1D | RPS10 | SSR3 | PTGDS | CCDC85B |
| CF | **FBLN1** | TIMP1 | SELENOM | LGALS1 | EIF1B | FTL | LRP1 | CKB | IFITM3 | IGFBP5 |
| EC | PTGDS | ENO1 | GAPDH | NFKBIA | IRF1 | ITM2C | NDUFA1 | TPI1 | ZFHX3 | IER3 |
| Ves | GNG11 | EGFL7 | **VIM** | TGFBR2 | TFPI | CRIP2 | CTSH | CD74 | ARHGAP29 | MGST2 |
| Mel | DCT | APOE | **PMEL** | **TYRP1** | GPX3 | **VIM** | QPCT | CD63 | CD59 | MT1G |
| IC | CYBA | B2M | STK4 | SRGN | ZFP36L2 | PTPRC | WIPF1 | CXCR4 | CREM | HSPD1 |
| MC | TAGLN | **MYL9** | PTMA | NR2F2 | DSTN | CALD1 | TPM1 | PPP1R12A | SEPTIN7 | IFITM3 |

D

| | gene 1 | gene 2 | gene 3 | gene 4 | gene 5 | gene 6 | gene 7 | gene 8 | gene 9 | gene 10 |
|---|---|---|---|---|---|---|---|---|---|---|
| LSC-1 | MT1X | DST | **SLC6A6** | NOP53 | RPS3A | RPS27 | PTMA | RPS2 | EZR | ID3 |
| LSC-2 | STMN1 | TKT | SAT1 | RPS4X | MT1X | RPL5 | SLC38A2 | ANP32B | RPS18 | RPL37A |
| LE | RPS10 | HMGA1 | GAPDH | FABP5 | GJA1 | S100A10 | HMGN3 | SLC25A6 | RPS8 | BZW1 |
| Cj | S100A11 | BTG1 | MGST1 | SAT1 | **CLDN4** | PRDX1 | FTH1 | KRT19 | TMSB4X | SLC2A1 |
| SK | RPS4X | HSPD1 | EIF5A | ANXA5 | **VIM** | EIF1AX | IFITM2 | HSP90AB1 | RPS27 | SLC2A1 |
| TSK | MT1G | MT1X | NOP53 | MT1E | SSR3 | DHX36 | EEF1D | NUDC | MT2A | CCNI |
| CF | **FBLN1** | TIMP1 | EIF1B | COL3A1 | FTL | **COL1A1** | AMD1 | NPM1 | COL1A2 | IFITM3 |
| EC | ENO1 | ZFHX3 | APOE | APP | NDUFA1 | TMEM160 | ATP5ME | FUS | EPB41L2 | UQCR11 |
| Ves | KDR | TFPI | MMRN1 | GNG11 | CALCRL | EGFL7 | LDB2 | **PECAM1** | ARHGAP29 | MEF2C |
| Mel | DCT | **PMEL** | APOE | EDNRB | CD63 | STMN1 | **VIM** | MYO5A | RBM3 | CHCHD6 |
| IC | HSPD1 | CYBA | ANKRD28 | B2M | ZEB2 | ZFP36L2 | DHX36 | AKAP13 | STK4 | LCP1 |
| nm-cSC | GPM6B | CALM2 | NRXN1 | SAMHD1 | WSB1 | KCNMB4 | WNT6 | HSP90AB1 | MT1G | CHPT1 |
| MC | PPP1R12A | SEPTIN7 | CALD1 | DSTN | TPM1 | NR2F2 | ID4 | PTMA | **NOTCH3** | **MYL9** |

E

| | gene 1 | gene 2 | gene 3 | gene 4 | gene 5 | gene 6 | gene 7 | gene 8 | gene 9 | gene 10 |
|---|---|---|---|---|---|---|---|---|---|---|
| LSC-1 | DST | **SLC6A6** | RPS3A | RPS27 | MT1X | NOP53 | RPL5 | RPL13 | BTF3 | RPL10 |
| LSC-2 | SAT1 | STMN1 | FTH1 | CREB5 | TKT | ANXA1 | CSRP2 | KCNMA1 | BTG1 | SLC38A2 |
| LE | GAPDH | RPS10 | RPS8 | SH3BGRL3 | KRT17 | TNFRSF12A | UBC | RPS9 | RPS27 | S100A10 |
| Cj | S100A11 | **CLDN4** | SAT1 | KRT19 | BTG1 | FTH1 | TMSB4X | CD55 | PERP | AQP3 |
| CE | CD24 | FTH1 | B2M | ELP4 | IGFBP2 | ELF3 | TKT | C4orf3 | CALM2 | RPL3 |
| SK | RPS4X | **VIM** | RPS8 | RPS18 | HSP90AB1 | ANXA5 | ANXA1 | RPS2 | HSPD1 | RPL10 |
| TSK | EEF1D | NOP53 | RPL13 | RPLP0 | CCNI | RPS10 | SSR3 | MTDH | AHI1 | EEF1A1 |
| CF | TIMP1 | **FBLN1** | IGFBP5 | FTL | RPS4X | RPS9 | EIF1B | CKB | DCT | HLA-E |
| EC | ENO1 | ZFHX3 | GAPDH | PTGDS | FTL | PCDH7 | APP | MIF | SFRP1 | NDUFA1 |
| Ves | ARHGAP29 | **VIM** | TFPI | TCF4 | FTH1 | EGFL7 | ZNF385D | MARCKSL1 | HLA-E | TMSB10 |
| Mel | DCT | **PMEL** | **TYRP1** | APOE | IGFBP5 | CD63 | TRPM1 | TIMP1 | **VIM** | MITF |
| IC | CXCR4 | PTPRC | B2M | CYBA | STK4 | ZEB2 | WIPF1 | ANKRD28 | FTH1 | EVI2B |
| nm-cSC | NRXN1 | GPM6B | CALM2 | **VIM** | SORBS2 | WSB1 | CHPT1 | HSP90AB1 | KCNMB4 | CST3 |
| MC | PPP1R12A | EBF1 | NR2F2 | SEPTIN7 | PTMA | CALD1 | DSTN | TPM1 | ZEB2 | LPP |

F

| | gene 1 | gene 2 | gene 3 | gene 4 | gene 5 | gene 6 | gene 7 | gene 8 | gene 9 | gene 10 |
|---|---|---|---|---|---|---|---|---|---|---|
| LSC-1 | MT1X | DST | NOP53 | RPS27 | RPS3A | **SLC6A6** | TXNIP | WFDC2 | BTF3 | ID3 |
| LSC-2 | ANXA1 | STMN1 | FTH1 | TKT | ALDH3A1 | SAT1 | **KRT14** | GLUL | ANP32B | NPM1 |
| LE | RPS10 | GAPDH | RPS8 | TNFRSF12A | SH3BGRL3 | CST3 | S100A10 | HMGN3 | SFN | SLC25A6 |
| Cj | S100A11 | BTG1 | **CLDN4** | KRT19 | SAT1 | MT1X | FTH1 | VEGFA | BAG1 | PERP |
| CE | CD24 | S100A11 | MT1X | TKT | S100A14 | **CLDN4** | KRT17 | PERP | ELF3 | **LYPD2** |
| TSK | MT1X | NOP53 | EEF1D | CCNI | SSR3 | MTDH | AHI1 | RPLP0 | NUDC | RPS10 |
| CF | TIMP1 | **FBLN1** | IGFBP5 | FTL | EIF1B | CKB | PLIN2 | **VIM** | SELENOM | ZFP36L2 |
| EC | PTGDS | ENO1 | ZFHX3 | GAPDH | PCDH7 | APP | TSPYL2 | DDB1 | MIF | NDUFA1 |
| Ves | TFPI | EGFL7 | ARHGAP29 | TGFBR2 | HYAL2 | MGST2 | TCF4 | CD59 | **VIM** | MARCKSL1 |
| Mel | DCT | **PMEL** | **TYRP1** | APOE | CD63 | CD59 | QPCT | **VIM** | SDCBP | CHCHD6 |
| IC | EVI2B | PTPRC | CYBA | B2M | STK4 | STK17B | HLA-B | LCP1 | ZFP36L2 | HLA-C |
| nm-cSC | GPM6B | NRXN1 | **VIM** | CALM2 | CST3 | KCNMB4 | WSB1 | CHPT1 | SORBS2 | HSP90AB1 |
| MC | PPP1R12A | SEPTIN7 | DSTN | PTMA | CALD1 | NR2F2 | ID4 | **MYL9** | TIMP3 | TPM1 |

***Supplemental Figure 5: Top positively contributing explainable AI genes with SHAP analysis. A-F)*** *Top 10 explainable genes positively contributing to model decisions for Maiti data (A-B): from adult cornea (A) and from 4-month organoids (B), and for Swarup data (C-F): 4-month organoids (C), 1-month organoids (D), 2-month organoids (E), 3-month organoids (F). Adult markers for cell states are highlighted in bold.*

##### 1.2 Supplemental Table S1

| Primary antibodies | Host | Company | Catalogue Number | Dilution |
|---|---|---|---|---|
| CPVL | Rabbit polyclonal | Thermo-Fisher | PA5-63308 | 1:100 |
| SLC6A6 | Rabbit polyclonal | Fisher Scientific | 16500992 | 1:100 |
| p63 (ΔNp63) | Mouse monoclonal [4A4] | Abcam | ab735 | 1:100 |
| Keratocan | Rabbit polyclonal | Sigma-Aldrich | HPA039321 | 1:100 |
| Fibulin-1 | Rabbit polyclonal | Fisher Scientific | 16620295 | 1:100 |
| POU3F3 | Rabbit polyclonal | Abcam | ab247159 | 1:100 |
| TNNC1 | Mouse polyclonal [4C2] | Abcam | ab10231 | 1:100 |

**Supplemental Table S2 provided separately as an Excel file**
